# Design principles to tailor Hsp104 therapeutics

**DOI:** 10.1101/2024.04.26.591398

**Authors:** JiaBei Lin, Peter J. Carman, Craig W. Gambogi, Nathan M. Kendsersky, Edward Chuang, Stephanie N. Gates, Adam L. Yokom, Alexandrea N. Rizo, Daniel R. Southworth, James Shorter

## Abstract

The hexameric AAA+ disaggregase, Hsp104, collaborates with Hsp70 and Hsp40 via its autoregulatory middle domain (MD) to solubilize aggregated protein conformers. However, how ATP- or ADP-specific MD configurations regulate Hsp104 hexamers remains poorly understood. Here, we define an ATP-specific network of interprotomer contacts between nucleotide-binding domain 1 (NBD1) and MD helix L1, which tunes Hsp70 collaboration. Manipulating this network can: (a) reduce Hsp70 collaboration without enhancing activity; (b) generate Hsp104 hypomorphs that collaborate selectively with class B Hsp40s; (c) produce Hsp70-independent potentiated variants; or (d) create species barriers between Hsp104 and Hsp70. Conversely, ADP-specific intraprotomer contacts between MD helix L2 and NBD1 restrict activity, and their perturbation frequently potentiates Hsp104. Importantly, adjusting the NBD1:MD helix L1 rheostat via rational design enables finely tuned collaboration with Hsp70 to safely potentiate Hsp104, minimize off-target toxicity, and counteract FUS proteinopathy in human cells. Thus, we establish important design principles to tailor Hsp104 therapeutics.

**Graphical Abstract:** 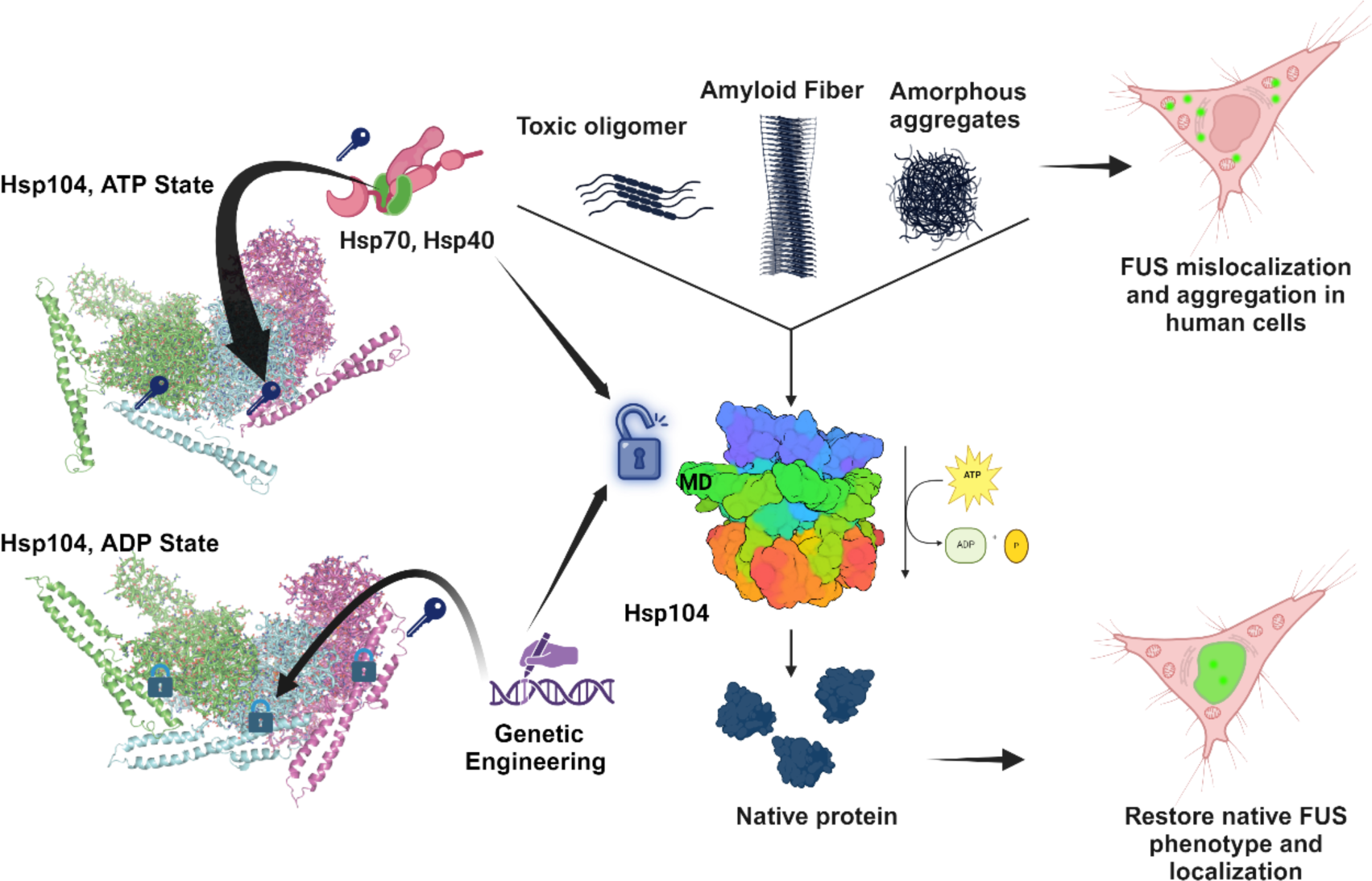

## Introduction

Protein function requires high-fidelity protein folding.^1^ However, in the cell, proteins are exposed to various stresses such as translational errors, heat or chemical shock, and aging, which can elicit protein misfolding and aggregation.^2^ The accumulation of misfolded and aggregated proteins is problematic and can be toxic.^3,4^ Thus, cells possess sophisticated molecular chaperones, protein disaggregases, protein-degradation systems, and stress-response pathways to maintain protein quality control.^5,6^ However, chronic accumulation of misfolded protein conformers upon aging can yield aberrant protein fibrils that are intimately tied to fatal neurodegenerative proteinopathies, such as α-synuclein fibrils in Parkinson’s disease (PD), or TDP-43 and FUS fibrils in amyotrophic lateral sclerosis (ALS).^7-10^ There are no effective treatments for these devastating neurodegenerative disorders.

One strategy for these neurodegenerative diseases would be to develop therapeutic protein disaggregases that liberate proteins trapped in aberrant oligomeric and aggregated states and restore them to native solubility, form, and function.^11^ Such agents would eliminate two malicious problems associated with deleterious protein misfolding and aggregation: (1) the toxic gain of function of aggregated conformers; and (2) the loss of protein function due to sequestration in aggregated conformers.^11^ Thus, we have focused on Hsp104, a ring-shaped, hexameric AAA+ (ATPase Associated with diverse Activities) protein disaggregase, which can suppress age-related protein aggregation.^11-14^ Notably, Hsp104 dissolves a diverse spectrum of aggregated structures, including preamyloid oligomers, disordered aggregates, phase-separated condensates, and stable amyloid or prion fibrils.^15-20^ However, no exact Hsp104 homolog is found in metazoa,^11,21,22^ although a related human mitochondrial AAA+ protein, Skd3, displays potent disaggregase activity,^23-26^ and another AAA+ protein VCP/p97 may remodel ubiquitylated protein inclusions in the cytoplasm.^27^ Nonetheless, introduction of Hsp104 or engineered variants into metazoan systems is well tolerated and can antagonize aggregation and toxicity of neurodegenerative disease proteins.^16,28-35^

Hsp104 is composed of two nucleotide-binding domains (NBD1 and NBD2) per monomer separated by a middle domain (MD) and flanked by an N-terminal domain (NTD) and C-terminal domain (CTD).^13^ The MD is an autoinhibitory domain that regulates Hsp104 disaggregase activity.^13^ Indeed, single mutations in the MD can relieve autoinhibition and enhance Hsp104 disaggregase activity.^36-41^ Precisely how the MD permits or restricts Hsp104 disaggregase activity is not completely understood.

Hsp70 and Hsp40 enable optimal Hsp104 disaggregase activity.^15,19,42-44^ Hsp70s, and their obligate cochaperones, Hsp40s, are highly conserved.^45^ Hsp40 binds substrate proteins and transfers them to Hsp70 via activation of Hsp70 ATPase activity.^46-48^ Interactions between Hsp70 and the NTD and MD of Hsp104 appear to enable disaggregase activity.^13,49-51^ However, the precise mechanism by which Hsp104, Hsp70, and Hsp40 coordinate activity remains uncertain. Several studies have utilized prokaryotic homologues of Hsp104 and Hsp70, ClpB and DnaK, respectively.^52-55^ However, prokaryotic ClpB is unable to perform the complete repertoire of eukaryotic Hsp104 activities.^17,18,31,50,56,57^ Indeed, there are several key structural and mechanistic differences between Hsp104 and ClpB.^50,57-61^ Several interaction sites on the MD of ClpB are proposed to interact with DnaK^52,53^ and the MD of Hsp104 has been proposed to interact with a fragment of human Hsp70 (HSPA1A).^49^ However, the structural determination of the Hsp104- Hsp70 interaction has been difficult to resolve due to the weak and transient interactions between these two proteins.^52,53^

We have discovered many Hsp104 variants bearing single missense mutations in NBD1, MD, or NBD2, which display enhanced disaggregase activity.^29,30,36,58,62-65^ However, some of these Hsp104 variants, particularly MD or NBD1 variants, can present with “off-target” toxicity.^29,30,63^ Specifically, overexpressing these Hsp104 variants in *Δhsp104* yeast reduces growth at 37°C, likely by unfolding metastable, soluble proteins.^29,36^ It has been suggested that Hsp70 might direct Hsp104 to aggregated proteins rather than misfolded soluble substrates, which may prevent off-target toxicity.^66^ However, the mechanism of Hsp104-Hsp70 cooperation and its connection to off-target toxicity is still poorly understood. Consequently, rational design of potentiated Hsp104 variants with no off-target toxicity remains a significant challenge.^30^

Here, we address these challenges by exploring an intimate network of contacts between the MD and NBD1 revealed in high-resolution cryogenic electron microscopy (cryo-EM) structures of Hsp104.^59^ We discover that the ATP-specific interactions between MD helix L1 and NBD1 of the adjacent clockwise protomer are critical for collaboration between Hsp104 and Hsp70 in protein disaggregation. Manipulating this network can: (a) reduce Hsp70 collaboration without enhancing activity; (b) generate Hsp104 hypomorphs that collaborate selectively with class B Hsp40s; (c) produce Hsp70-independent potentiated variants; or (d) create species barriers between Hsp104 and Hsp70. By contrast, the distinctive ADP-specific intraprotomer contacts between MD helix L2 and NBD1 restrict activity, and their perturbation frequently potentiates disaggregase activity. We establish that the off-target toxicity of specific potentiated Hsp104 variants is determined by reduced dependence on Hsp70 for protein disaggregation. By tuning the ATP-specific MD helix L1 and NBD1 interaction, we can specify a desired level of Hsp70 collaboration to yield potentiated Hsp104 variants with no off-target toxicity. Importantly, for the first time, we establish that potentiated Hsp104 variants can mitigate FUS proteinopathy in human cells. Overall, our findings establish important design principles to tailor therapeutic Hsp104 variants.

## Results

### The MD changes orientation as Hsp104 hexamers switch from ATP-bound to ADP-bound states, which alter NBD1:MD interactions

The MD plays a critical role in regulating Hsp104 activity.^13,29,67-69^ High-resolution structures of Hsp104 hexamers determined by cryo-EM reveal a dramatic change in the orientation of the MD between the substrate-free AMP-PNP state (hereafter referred to as the ATP state) and substrate-free ADP state (Figure 1A, B).^59,70^ Importantly, these two distinct states are also observed in substrate-bound Hsp104 hexamers in the corresponding protomers bound to ATP or ADP, indicating that these states are populated by distinct protomers during substrate translocation.^59^ However, it has remained unclear precisely how Hsp104 activity is regulated by the distinct interactions between NBD1 and the MD in either state.

**Figure 1.**
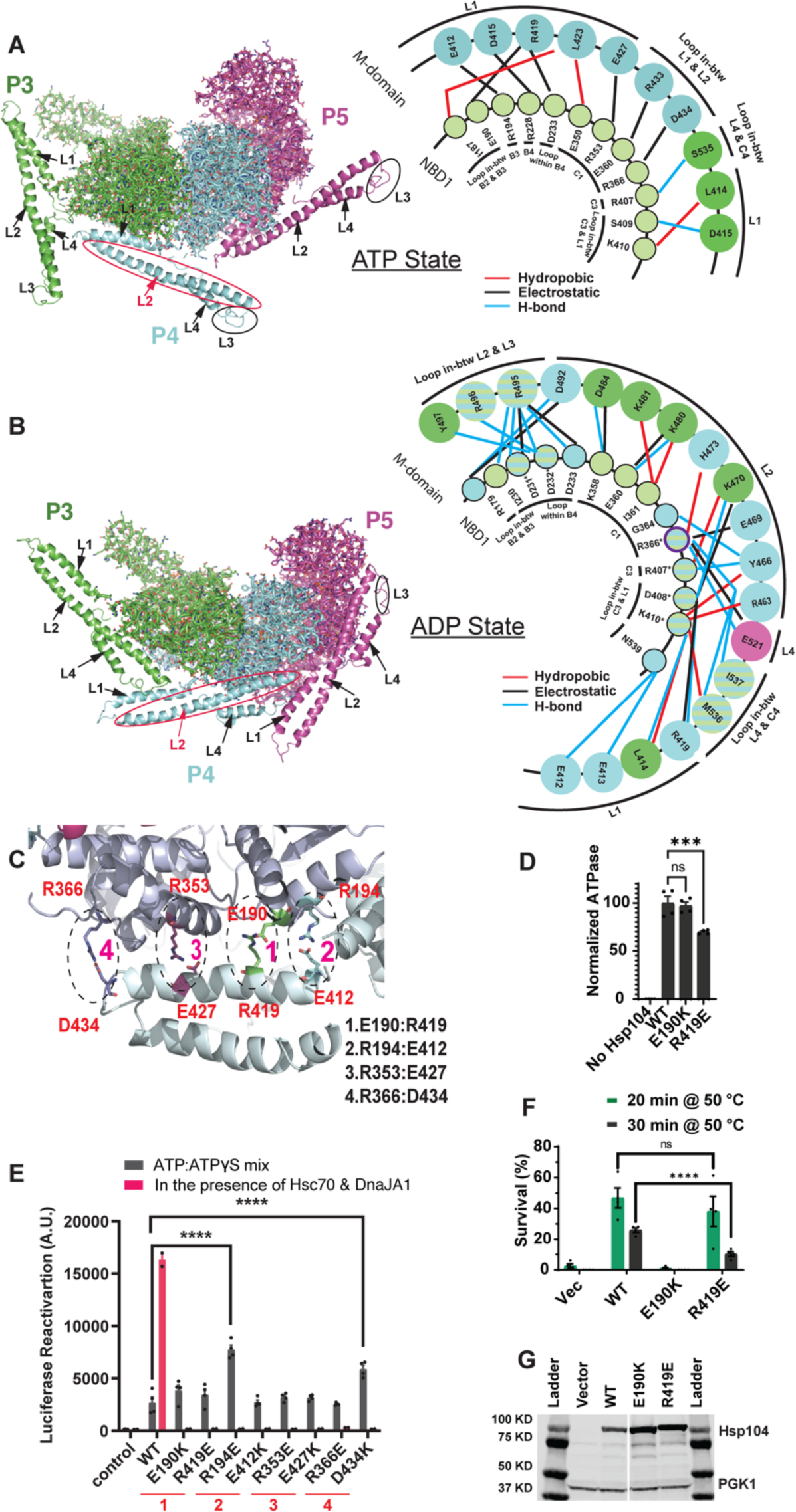
The MD changes orientation as Hsp104 hexamers switch from ATP-bound to ADP-bound states, which alter NBD1:MD interactions. **(A, B)** Left, three out of six protomers (P3 in green, P4 in blue, and P5 in magenta) are shown for the ATP state (A) or ADP state (B). NBD1 is shown in ribbon and MD is shown in cartoon. The four MD helices (L1, L2, L3, and L4) are indicated by arrows. However, MD helix L3 appears to be a loop in these structures. Right, the NBD1:MD interactions for protomers P3 (Green) and P4 (Blue) were analyzed using Discovery Studio Visualizer with a 4Å cut off distance for the ATP-bound state (AMP-PNP, 5KNE) (A) or the ADP-bound state (5VY8) (B). Hydrophobic, electrostatic (salt-bridge), and hydrogen bond interactions are indicated in red, black, and blue lines, respectively. For the ATP-bound state (A), NBD1:MD interactions were observed between the NBD1 of P3 (green circles) and MD of P4 (blue circles). For the ADP-bound state (B), we show NBD1:MD interactions within P3 (green circles) or within P4 (blue circles) and a unique interaction between R366 and E521 within P5 (purple) for clarity. Intrasubunit NBD1:MD interactions conserved in P3 and P4 are shown in circles with blue and green stripes. Residues involved in intrasubunit NBD1:MD interactions for P3, P4 or P5 are shown in circles with blue and green stripes with an ‘*’. **(C)** NBD1:MD helix L1 interactions with NBD1 in the ATP state. Four major salt-bridge interactions: 1. E190:R419, 2. R194:E412, 3. R353:E427 and 4. R366:D434, are identified on the inter-subunit NBD1:MD interface in the presence of AMP-PNP. E190, R194, R353, and R366 are in NBD1 of subunit P3, and R419, E412, E427 and D434 are in MD helix L1 of P4. **(D)** ATPase activity of Hsp104, Hsp104^E190K^, and Hsp104^R419E^ (0.25µM) in the presence of 1mM ATP at 25°C. Bars represent means±SEM (n=4); each replicate is shown as a dot. Ordinary one-way ANOVA Dunnett’s test was performed to compare the ATPase activity of Hsp104 to Hsp104^E190K^ or Hsp104^R419E^. ns=not significant; ***p=0.0005. **(E)** Hsp104 variants (1µM, monomer) in the presence of ATP:ATPγS (2.5mM:2.5 mM; black bars) or with Hsc70 (0.167µM) and DnaJA1 (0.167µM; pink bars) were incubated with 100nM (monomeric concentration) chemical-denatured luciferase aggregates for 90min at 25°C. Buffer serves as the negative control. Bars represent means±SEM (n=4); each replicate is shown as a dot. ****P≤0.0001. **(F)** Survival (%) of Δ*hsp104* yeast transformed with empty vector (pRS313HSE), Hsp104, Hsp104^E190K^, or Hsp104^R419E^ after 0, 20, or 30min heat shock at 50°C following a pretreatment at 37°C for 30min. Bars represent means±SEM (n=4); each replicate is shown as a dot. ns=not significant; ****P≤0.0001. **(G)** *Δhsp104* yeast from (F) were incubated at 37°C for 30 min to induce Hsp104 expression. Yeast were then lysed and processed for Western blot. 3-Phosphoglycerate kinase 1 (PGK1) serves as a loading control. See also Figure S1.

We surveyed the interactions between NBD1 and the MD for each nucleotide state (Figure 1A, B). Numerous contacts, including hydrophobic (red line), electrostatic (black line), and hydrogen bond (blue line) interactions, are observed in both nucleotide states (Figure 1A, B, right). All the interactions observed in protomer 3 (P3, in green) and protomer 4 (P4, in blue) are displayed for the ATP state (Figure 1A, right) and ADP state (Figure 1B, right). Notably, in the ATP state we find that helix L1 of the MD of protomer 4 makes several interprotomer contacts with NBD1 of the adjacent clockwise protomer 3 (Figure 1A).^70^

By contrast, in the ADP state, we find that interactions between NBD1 and the MD are completely remodeled such that helix L2 of the MD now makes intraprotomer contacts with NBD1 (Figure 1B).^59^ Moreover, in the ADP state, the precise NBD1:MD intraprotomer interactions are not completely identical in different subunits. Some interactions are only present in P3 (shown in green) or P4 (blue) (Figure 1B, right). Only a few interactions are the same in both protomers (Figure 1B, shown in green and blue stripes), indicating diverse intraprotomer interactions between the MD and NBD1 in this state. In addition to helix L2, helices L1 and L4 as well as the loops between helix L2 and L3 and helix L4 and NBD1 also make intraprotomer contacts with NBD1 (Figure 1B). Thus, the MD makes a radically different set of interactions with NBD1 in the presence of ADP versus ATP.

We hypothesized that the interprotomer interactions between NBD1 and the MD in the ATP state (Figure 1A) and intraprotomer interactions between NBD1 and the MD in the ADP state (Figure 1B) may play key roles in regulating Hsp104 activity. Thus, to investigate the mechanism by which the MD regulates Hsp104 activity, we performed mutagenesis analysis to perturb the interactions between NBD1 and the MD for each state. Remarkably, we find that perturbation of the interprotomer NBD1-MD interactions of the ATP state has different functional consequences for Hsp104 activity than perturbation of the intraprotomer NBD1-MD interaction of the ADP state.

### Interprotomer NBD1:MD interactions of the ATP state are essential for Hsp104 collaboration with Hsp70

To alter interactions between NBD1 and MD helix L1 in the ATP state, we first modified the charges of four of the NBD1:MD salt-bridge interactions: E190:R419, R194:E412, R353:E427, and R366:D434 (Figure 1C). Thus, we constructed the opposite charge variants to perturb the salt bridges: Hsp104^E190K^, Hsp104^R419E^, Hsp104^R194E^, Hsp104^E412K^, Hsp104^R353E^, Hsp104^E427K^, Hsp104^R366E^, and Hsp104^D434K^. Previously, we purified and measured the ATPase activity of Hsp104^R194E^, Hsp104^E412K^, Hsp104^R353E^, Hsp104^E427K^, Hsp104^R366E^, and Hsp104^D434K^.^59^ We found that these Hsp104 variants display wild-type (WT) levels of ATPase activity, which is stimulated by the disordered substrate casein.^59^ However, these variants cannot work with human Hsp70 (Hsc70 [HSPA8]) and Hsp40 (DnaJA1) to disaggregate and reactivate firefly luciferase trapped in chemically denatured aggregates.^59^ We have now purified the remaining variants, Hsp104^E190K^ and Hsp104^R419E^, which possessed WT levels (Hsp104^E190K^) or approaching WT levels (∼70% for Hsp104^R419E^) of ATPase activity (Figure 1D). Hsp104^E190K^ and Hsp104^R419E^ also displayed limited ability to work with human Hsc70 and DnaJA1 to disaggregate and reactivate luciferase (Figure 1E, pink bars). These findings suggest that perturbation of the MD helix L1 interactions with NBD1 of the adjacent clockwise protomer in the ATP state does not grossly affect ATPase activity but reduces disaggregase activity.

It remained unclear, however, whether disruption of these contacts prevents Hsp104 from coupling ATP hydrolysis to protein disaggregation or whether they specifically reduce collaboration with Hsp70 and Hsp40. To distinguish between these possibilities, we first established that these Hsp104 variants bind a model, disordered substrate, β-casein, with the same affinity as Hsp104 (Figure S1A). Thus, substrate engagement appears unaffected by these mutations. We next assessed the intrinsic disaggregase activity of the Hsp104 variants in the presence of a 1:1 ratio of ATP and the slowly hydrolyzable ATP analog, ATPγS. Hsp104 disaggregates and reactivates luciferase trapped in chemically denatured aggregates in the presence of a 1:1 ratio of ATP:ATPγS in the absence of Hsp70 and Hsp40,^18,65,71^ thereby allowing assessment of whether the mutations perturbed the ability of Hsp104 to couple ATP hydrolysis to protein disaggregation. We found that all the Hsp104 variants tested could disaggregate and reactivate luciferase as well or better than WT Hsp104 under these conditions (Figure 1E, grey bars). Indeed, Hsp104^R194E^ and Hsp104^D434K^ were more effective than Hsp104 (Figure 1E, grey bars). However, unlike Hsp104, none of these Hsp104 variants could collaborate with human Hsc70 and DnaJA1 to disaggregate and reactivate luciferase (Figure 1E, pink bars). These findings suggest that perturbation of the MD helix L1 interactions with NBD1 of the adjacent clockwise protomer in the ATP state can specifically impair Hsp70 collaboration and do not affect the ability of Hsp104 to couple ATP hydrolysis to protein disaggregation.

We next tested the ability of these Hsp104 variants to confer induced thermotolerance (i.e., ability to survive at 50°C after a 37°C pretreatment) in Δ*hsp104* yeast. Previously, we tested Hsp104^R194E^, Hsp104^E412K^, Hsp104^R353E^, Hsp104^E427K^, Hsp104^R366E^, or Hsp104^D434K^, which were unable to confer induced thermotolerance.^59^ Hsp104^R366E^ displayed some limited activity, which was greater than the other variants, but was still largely ineffective.^59^ These findings suggest that Hsp104 collaboration with Hsp70 and Hsp40 is critical for thermotolerance *in vivo*.^72^ We now extend this analysis to Hsp104^E190K^ and Hsp104^R419E^. Hsp104^E190K^ exhibited impaired activity as with other NBD1:MD variants^59^ (Figure 1F) and was expressed at similar levels to Hsp104 (Figure 1G). Surprisingly, however, Hsp104^R419E^ could confer induced thermotolerance like Hsp104 after 20min but displayed reduced activity at 30min (Figure 1F). Hsp104, Hsp104^E190K^, and Hsp104^R419E^ were all expressed at similar levels (Figure 1G). One possible explanation for the activity of Hsp104^R419E^ and the residual activity of Hsp104^R366E^ might be an ability to collaborate with a subset of Hsp70 or Hsp40 homologues in yeast.

### Residues and salt-bridge interactions in the interprotomer NBD1:MD helix L1 interface regulate Hsp40 compatibility for Hsp104-Hsp70 disaggregase activity

To assess this possibility, we purified yeast Hsp70 homologue, Ssa1, class A Hsp40, Ydj1, and class B Hsp40, Sis1, to test their ability to collaborate with the Hsp104 variants in luciferase disaggregation and reactivation *in vitro*. Hsp104^E190K^, Hsp104^R194E^, Hsp104^E412K^, Hsp104^R353E^, Hsp104^E427K^, and Hsp104^D434K^ cannot disaggregate and refold luciferase in the presence of Ssa1 plus Sis1, Ssa1 plus Ydj1, or Ssa1 plus Sis1 and Ydj1 (Figure 2A). Thus, these Hsp104 variants are severely impaired in collaboration with Ssa1, Sis1, and Ydj1. By contrast, Hsp104^R419E^ and Hsp104^R366E^ displayed some activity in the presence of Ssa1 plus Sis1 or Ssa1 plus Sis1 and Ydj1, but not Ssa1 plus Ydj1 (Figure 2A). Thus, Hsp104^R419E^ and Hsp104^R366E^ are selectively defective in collaboration with the class A Hsp40, Ydj1.

**Figure 2.**
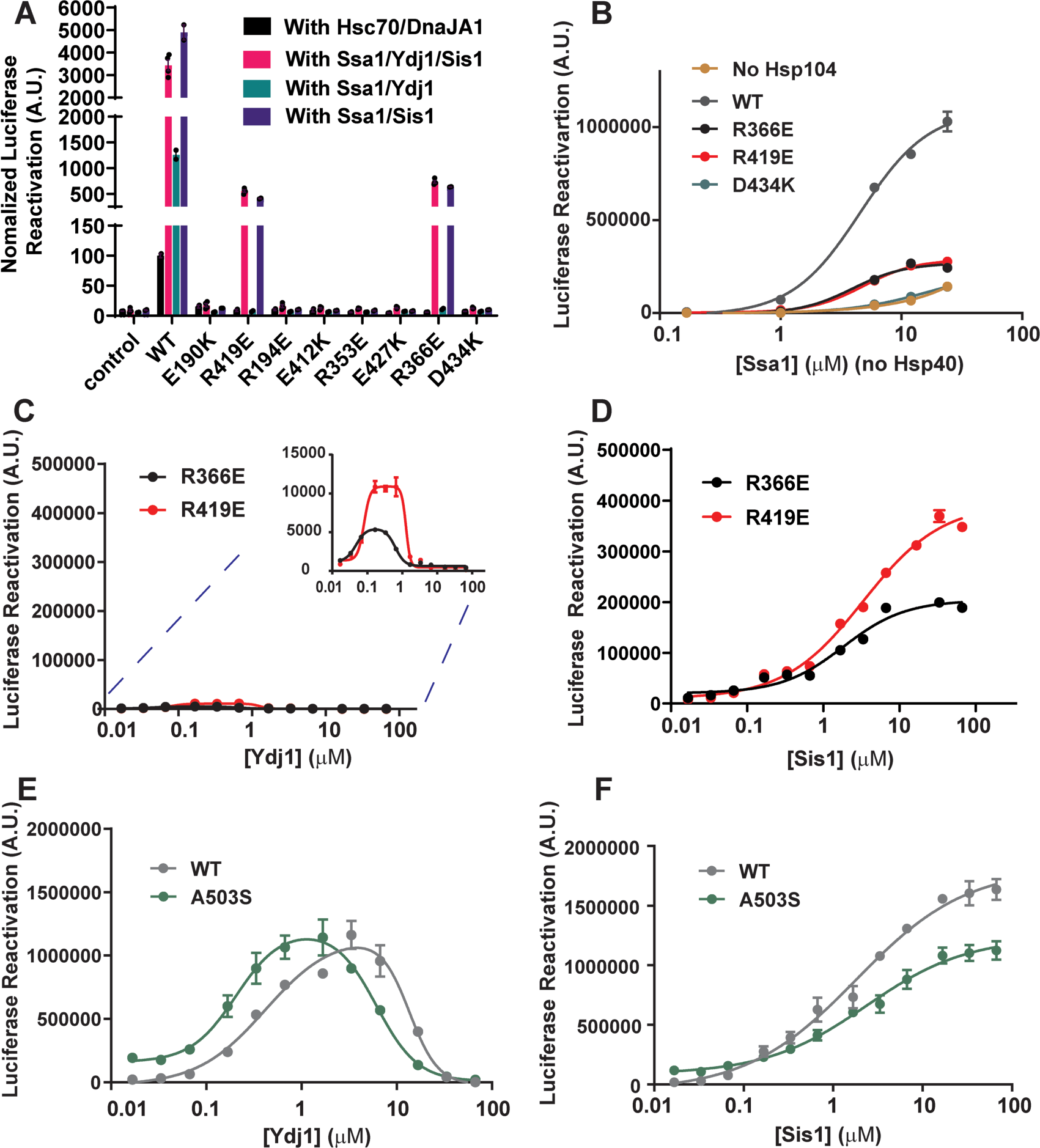
Specific perturbation of ATP-specific NBD1:MD helix L1 contacts yields Hsp104 variants that collaborate selectively with class B Hsp40s. **(A)** Luciferase disaggregation and reactivation activity of the indicated Hsp104 variants (1µM, monomeric) in the presence of Hsc70 (0.167µM) and DnaJA1 (0.167µM; black), Ssa1 (0.167µM), Ydj1 (0.083µM), and Sis1 (0.083µM; pink), Ssa1 (0.167µM) and Ydj1 (0.167µM; green), or Ssa1 (0.167µM) and Sis1 (0.167µM; purple). Bars represent means±SEM (n=2-4), each replicate is shown as a dot. **(B)** Luciferase disaggregation and reactivation activity of Hsp104 (grey dots), Hsp104^R366E^ (black dots), Hsp104^R419E^ (red dots), or Hsp104^D434K^ (teal dots; 1µM, monomeric) in the presence of various Ssa1 (Hsp70) concentrations in the absence of Hsp40. Curves were fit for absolute EC_50_ (see STAR Methods and Table S1). Values represent means±SEM (n=2). **(C, D)** Luciferase disaggregation and reactivation activity of Hsp104^R366E^ (1µM monomeric) plus Ssa1 (0.167µM; black) or Hsp104^R419E^ (1µM monomeric) plus Ssa1 (0.167µM; red) as a function of Ydj1 (C) or Sis1 (D) concentration. Curves were fit for IC_50_ of Ydj1 and EC_50_ of Ydj1 or Sis1 (see STAR Methods and Table S1). Values represent means±SEM (n=2). **(E, F)** Luciferase disaggregation and reactivation activity of Hsp104 (1µM, monomeric) plus Ssa1 (0.167µM; grey) or Hsp104^A503S^ (1µM, monomeric) plus Ssa1 (0.167µM; green) as a function of Ydj1 (E) or Sis1 (F) concentration. Curves were fit for IC_50_ of Ydj1 and EC_50_ of Ydj1 or Sis1 (see STAR Methods and Table S1). Values represent means±SEM (n=2). See also Figure S1, S2 and Table S1.

We next asked whether Hsp104^R419E^ and Hsp104^R366E^ collaboration with Hsp70 was also impaired. Thus, we tested the Ssa1 dose-dependent disaggregation activity of Hsp104 (positive control), Hsp104^R366E^, Hsp104^R419E^, and Hsp104^D434K^ (negative control) in the absence of Hsp40. Hsp104^D434K^ is incapable of working with Ssa1 to disaggregate luciferase, whereas Hsp104^R419E^ and Hsp104^R366E^ remain partially active, but have reduced activity compared to Hsp104 (Figure 2B). Together with the yeast thermotolerance results (Figure 1F),^59^ our findings suggest that Hsp104^E190K^, Hsp104^R194E^, Hsp104^E412K^, Hsp104^R353E^, Hsp104^E427K^, and Hsp104^D434K^ are completely impaired for collaboration with Hsp70, whereas Hsp104^R419E^ and Hsp104^R366E^ can partially work with Hsp70.

Next, we examined Hsp40 compatibility with Hsp104^R419E^ and Hsp104^R366E^ in more detail. Thus, we assessed luciferase disaggregation and reactivation at fixed Hsp104^R419E^ or Hsp104^R366E^ (1µM) and Ssa1 (0.167µM) concentrations with increasing amounts of Sis1 or Ydj1. Remarkably, Hsp104^R419E^ and Hsp104^R366E^ were unable to collaborate with Ydj1 (Figure 2C), whereas they could collaborate with Sis1 (Figure 2D). By contrast, Hsp104 and a canonical potentiated variant, Hsp104^A503S^,^29^ were active with Ydj1 or Sis1, although Ydj1 inhibited activity at high concentrations, whereas Sis1 did not (Figure 2E, F). Notably, Hsp104^A503S^ is much more active than Hsp104 at low Ydj1 concentrations (Figure S1B).^29^ However, at the optimal Sis1 or Ydj1 concentrations, the disaggregase activities of Hsp104 and Hsp104^A503S^ are similar (Figure 2E, F). Our findings suggest that Hsp104^R419E^ and Hsp104^R366E^ can partially work with Ssa1 (Figure 2B), but their luciferase reactivation activity is more sensitive to inhibition by Ydj1 (Figure 2C). Thus, Hsp104 collaboration with Ydj1 during luciferase disaggregation and reactivation requires R366 in NBD1 and R419 in MD helix L1.

### Ydj1 but not Sis1 can dissociate substrates from Hsp104 and inhibit the spontaneous refolding of unfolded luciferase

Bell-shaped reactivation curves were observed for all Hsp104 variants as a function of Ydj1 concentration, indicating that the synergistic cooperativity of Hsp104-Ssa1-Ydj1 is regulated by the Ydj1 concentration (Figure 2C, E). In the tested concentration range, an inhibitory effect of Sis1 on luciferase disaggregation and reactivation activity was not observed (Figure 2D, F). The inhibition of protein disaggregation and reactivation by high Ydj1 concentrations could be due to substrate competition between Ydj1 and Hsp104-Hsp70 complex, which may resemble the ‘hook effect’ of proteolysis targeting chimera (PROTAC) molecules (i.e., high concentrations of a linker [in this case Ydj1] suppresses formation of ternary complexes due to excessive formation of binary complexes).^73^ Additionally or alternatively, this effect might be explained by excess Ydj1 binding to unfolded luciferase released by Hsp104 and preventing it from refolding.

The class A Hsp40, Ydj1, has six sites involved with substrate binding in each dimer.^74^ These sites reside in client-binding domain 1 (CBD1), client-binding domain 2 (CBD2), and the zinc-finger domain.^74^ By contrast, the class B yeast Hsp40, Sis1, contains CBD1, but lacks CBD2.^74^ Thus, Ydj1 may have a higher affinity for interacting with the substrate than Sis1, which might compete with Hsp104 for substrate binding. To test this idea, we preformed Hsp104:β-casein complexes in the presence of ATPγS and titrated in Ydj1 or Sis1. Hsp104^R419E^ and Hsp104^R366E^ bind to casein with similar affinity as Hsp104 (Figure S1). As predicted, Ydj1 can compete for β-casein (30nM) from Hsp104^R419E^, Hsp104^R366E^, and Hsp104 at an IC_50_∼20-40μM (Figure S2A, Table S1), whereas Sis1 cannot (Figure S2B). This IC_50_ value is similar to the IC_50_ of Ydj1 for inhibition of luciferase disaggregation and reactivation by Hsp104 at ∼14μM (Figure 2C, Table S1). By contrast, Hsp104^R419E^ and Hsp104^R366E^ are inhibited by much lower concentrations of Ydj1 in luciferase disaggregation and reactivation (Figure 2C and Table S1). These results suggest that Ydj1 may compete for substrate binding to Hsp104, which may contribute to the inhibition of Hsp104-mediated luciferase disaggregation and reactivation at high Ydj1 concentrations. However, this mode of inhibition does not readily explain the inhibition of Hsp104^R419E^ and Hsp104^R366E^ by low Ydj1 concentrations.

We wondered whether Ydj1 might inhibit the refolding of the small amounts of luciferase released by Hsp104^R419E^ and Hsp104^R366E^, which would severely restrict their luciferase reactivation activity. Thus, we next tested whether Ydj1 or Sis1 can inhibit spontaneous refolding of soluble unfolded luciferase. For this purpose, we unfold native luciferase (at the luciferase concentrations indicated in Figure S2C) with 6M Urea on ice for 5min. We then dilute the soluble unfolded luciferase into buffer and verified the spontaneous refolding of luciferase over 90 min (Figure S2C). At high concentrations, Ydj1 can inhibit this spontaneous refolding of soluble luciferase, but this inhibition is dependent on the luciferase concentration (Figure S2D). For low concentrations of unfolded luciferase (1nM or 2nM), Ydj1 inhibits spontaneous refolding with an IC_50_ of ∼3μΜ (Figure S2D). At higher concentrations of unfolded luciferase (10nM or 20nM), inhibition by Ydj1 is insignificant, and the IC_50_ cannot be determined at the tested Yjd1 concentration range (Figure S2D). By contrast, Sis1 does not inhibit luciferase refolding at all tested luciferase concentrations (Figure S2E). These results indicate that when small amounts of luciferase are released from aggregates by Hsp104, its refolding can be inhibited by excess Ydj1. We suggest that Hsp104 releases more unfolded luciferase than Hsp104^R419E^ and Hsp104^R366E^, and thus higher Ydj1 concentrations are needed to inhibit Hsp104 than Hsp104^R419E^ or Hsp104^R366E^. Collectively, these findings suggest that reduced ability to collaborate with Ssa1 and inhibition by Ydj1 render Hsp104^R419E^ and Hsp104^R366E^ hypomorphic *in vitro* and *in vivo*. These findings also emphasize the importance of Hsp104 collaboration with Ssa1 and Ydj1 for thermotolerance *in vivo*, as the Hsp104 variants that can only work with Ssa1 and Sis1 are hypomorphic.^72,75,76^

### Rewiring the interprotomer NBD1:MD helix L1 interaction alters Hsp104 collaboration with Hsp70 and Hsp40

We next tested whether rewiring salt-bridge interactions between NBD1 and MD helix L1 could restore Hsp104 collaboration with Hsp70 and Hsp40. Thus, we generated Hsp104^E190K:R419E^, Hsp104^E190R:R419E^, Hsp104^R194E:E412K^, Hsp104^R353E:E427K^, Hsp104^R353E:E427R^, Hsp104^R366E:D434K^, and Hsp104^R366E:D434R^, which would be predicted to have reconstructed salt bridges between NBD1 and MD helix L1 in the ATP state (Figure 1C). However, in these variants the location of the acidic and basic partner residues is reversed compared to Hsp104.

We first assessed the activity of these Hsp104 variants in luciferase disaggregation and reactivation. To our surprise, Hsp104^E190K:R419E^ and Hsp104^E190R:R419E^ show enhanced activity compared to Hsp104 in the absence or presence of human Hsc70 and DnaJA1 (Figure 3A). Remarkably, Hsp104^E190R:R419E^ exhibited similar activity with or without Hsc70 and DnaJA1 (Figure 3A). The increased level of activity of Hsp104^E190K:R419E^ and Hsp104^E190R:R419E^ and their independence from Hsp70 is consistent with potentiated Hsp104 activity.^29,30,40,51,58,63,64^ Indeed, these variants also exhibited increased ATPase activity, another indicator of potentiated activity (Figure S3A).^29,30,40,51,58,63,64^ Thus, altering the charge orientation of the salt bridge between position 190 of NBD1 and position 419 of MD helix L1 potentiates Hsp104 activity (Figure 3A). Intriguingly, this potentiated activity was only uncovered at this position of the NBD1:MD helix L1 interaction, which may pinpoint a precise location where Hsp70 activates Hsp104.

**Figure 3.**
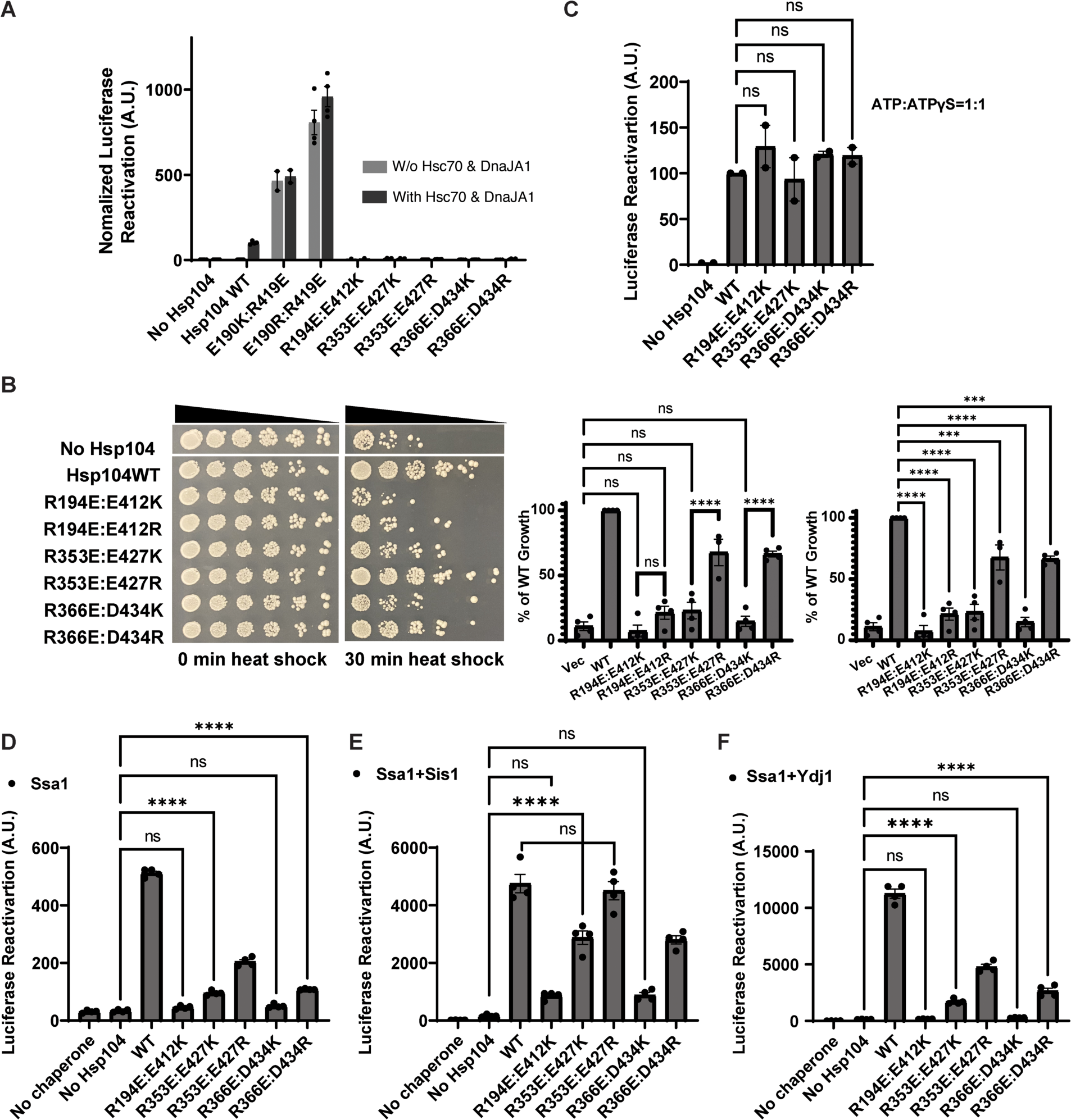
Rewiring the ATP-specific interprotomer NBD1:MD helix L1 interaction alters Hsp104 collaboration with Hsp70 and Hsp40. **(A)** Luciferase disaggregation and reactivation activity of the indicated Hsp104 variants (1µM, monomeric) in the absence (grey bars) or presence (black bars) of Hsc70 (0.167µM) and DnaJA1 (0.167µM). Bars represent means±SEM (N=2-4); each replicate is shown as a dot. One-way ANOVA Dunnett’s test was performed to compare Hsp104 to Hsp104 variants at a 95% confidence interval (CI). The luciferase disaggregation and reactivation activity of all tested Hsp104 variants is significantly different from Hsp104 with ****P≤0.0001 (statistics are omitted for clarity). **(B)** Left, spotting assay to evaluate the survival of Δ*hsp104* yeast transformed with empty vector (no Hsp104) or the indicated Hsp104 variants (WT=wild type) after a 30min pretreatment at 37°C followed by a 30min heat shock at 50°C. Δ*hsp104* yeast that were not heat shocked are shown on the left as a control. Right panels, yeast survival (% of wild-type Hsp104) was quantified. Bars represent means±SEM (n=4), and each replicate is shown as a dot. One-way ANOVA Dunnett’s tests were performed to compare Hsp104 variants to no Hsp104 control (middle panel) or WT (right panel) at 95% CI. ns=not significant; ***P≤0.001 ****P≤0.0001. One-way ANOVA Tukey tests were performed to make pairwise comparisons between specific Hsp104 variants as indicated (middle panel) ****P≤0.0001. **(C)** Luciferase disaggregation and reactivation activity of the indicated Hsp104 variants (1µM, monomeric) in the presence of ATP:ATPγS (2.5mM:2.5mM) and the absence of Hsp70 and Hsp40. Bars represent means±SEM (n=2), and each replicate is shown as a dot. One-way ANOVA Dunnett’s test was performed to compare Hsp104 variants to Hsp104 at 95% CI. ns=not significant. **(D)** Luciferase disaggregation and reactivation activity of the indicated Hsp104 variants (1µM monomeric) in the presence of Ssa1 (0.167µM) and absence of Hsp40. Bars represent means±SEM (n=4); each replicate is shown as a dot. One-way ANOVA Dunnett’s test was performed to compare Hsp104 variants to WT Hsp104 at 95% CI. All variants have significantly reduced activity to work with Ssa1 compared to WT Hsp104 with a P≤0.0001 (statistics are omitted for clarity). Alternatively, one-way ANOVA Dunnett’s test was performed to compare Hsp104 variants to no Hsp104. ns=not significant; ****P≤0.0001. **(E)** Luciferase disaggregation and reactivation activity of the indicated Hsp104 variants (1µM, monomeric) in the presence Ssa1 (0.167µM) and Sis1 (0,167µM). Bars represent means±SEM (n=4), and each replicate is shown as a dot. One-way ANOVA Dunnett’s test was performed to compare Hsp104 variants to WT Hsp104 at 95% CI. ns=not significant. All other variants have reduced activity with a P≤0.0001 (statistics are omitted for clarity). Alternatively, one-way ANOVA Dunnett’s test was performed to compare Hsp104 variants to no Hsp104. Ns=not significant, ****P≤0.0001. **(F)** Luciferase disaggregation and reactivation activity of the indicated Hsp104 variants (1µM, monomeric) in the presence of Ssa1 (0.167µM) and Ydj1 (0.167µM). Bars represent means±SEM (n=4); each replicate is shown as a dot. One-way ANOVA Dunnett’s test was performed to compare Hsp104 variants to WT Hsp104 at 95% CI. All Hsp104 variants have reduced activity with a ****P≤0.0001 (statistics are omitted for clarity). Alternatively, one-way ANOVA Dunnett’s test was performed to compare Hsp104 variants to no Hsp104. ns=not significant, ****P≤0.0001. See also Figure S3.

Indeed, Hsp104^R194E:E412K^, Hsp104^R353E:E427K^, Hsp104^R353E:E427R^, Hsp104^R366E:D434K^, and Hsp104^R366E:D434R^ were inactive for luciferase disaggregation and reactivation in the presence or absence of human Hsc70 and DnaJA1 (Figure 3A). These variants retained ATPase activity, but it was reduced in comparison to Hsp104 (Figure S3A). To determine activity *in vivo*, we assessed their ability to confer induced thermotolerance in Δ*hsp104* yeast. As expected, Hsp104^R194E:E412K^, Hsp104^R194E:E412R^, Hsp104^R353E:E427K^, and Hsp104^R366E:D434K^ were unable to confer induced thermotolerance (Figure 3B). To our surprise, however, Hsp104^R353E:E427R^ and Hsp104^R366E:D434R^ conferred induced thermotolerance to ∼67% of the level conferred by Hsp104 (Figure 3B). These Hsp104 variants were expressed at similar levels (Figure S3B). Thus, Hsp104^R353E:E427R^ and Hsp104^R366E:D434R^ are functional *in vivo* but fail to collaborate with human Hsc70 and DnaJA1 *in vitro* (Figure 3A, B).

To understand the concordance (for Hsp104^R194E:E412K^, Hsp104^R353E:E427K^, and Hsp104^R366E:D434K^) and discordance (for Hsp104^R353E:E427R^ and Hsp104^R366E:D434R^) between the *in vitro* and *in vivo* data, we further dissected the activity of these variants *in vitro*. First, we found that the intrinsic luciferase disaggregation and reactivation activity of Hsp104^R194E:E412K^, Hsp104^R353E:E427K^ and Hsp104^R366E:D434K^ in the presence of ATP:ATPγS and the absence of Hsp70 and Hsp40 was like Hsp104 (Figure 3C). Thus, these variants can couple ATP hydrolysis to protein disaggregation and reactivation, indicating a specific defect in collaboration with Hsp70 and Hsp40. These data reinforce the concept that the ATP-specific NBD1:MD helix L1 interaction is critical for collaboration with Hsp70 and Hsp40.

Next, we assessed whether the functional interaction with Ssa1 was affected. Thus, we assessed luciferase disaggregation and reactivation in the presence of ATP plus Ssa1, but in the absence of Hsp40. Here, none of the variants had levels of activity comparable to Hsp104 (Figure 3D). Indeed, Hsp104^R194E:E412K^ and Hsp104^R366E:D434K^ were inactive, whereas Hsp104^R353E:E427K^ and Hsp104^R366E:D434R^ exhibited limited activity (Figure 3D). Hsp104^R353E:E427R^ exhibited ∼40% Hsp104 activity, which is consistent with the ability of this variant to confer some induced thermotolerance (Figure 3B). Nonetheless, these findings suggest that these Hsp104 variants have reduced ability to collaborate directly with Ssa1.

We then added the class B Hsp40, Sis1, or class A Hsp40, Ydj1, together with Ssa1 and assessed luciferase disaggregation and reactivation activity (Figure 3E, F). Here, Hsp104^R194E:E412K^ and Hsp104^R366E:D434K^ are inactive with Ssa1 plus Sis1 or Ydj1 (Figure 3E, F), which explains the inability of these Hsp104 variants to confer induced thermotolerance (Figure 3B). Hsp104^R353E:E427K^ and Hsp104^R366E:D434R^ both exhibited ∼60% of Hsp104 activity with Ssa1 plus Sis1 (Figure 3E), and limited activity with Ssa1 and Ydj1 (Figure 3F). Hsp104^R366E:D434R^ (∼24% of Hsp104) was slightly more active with Ssa1 and Ydj1 than Hsp104^R353E:E427K^ (∼15% of Hsp104) (Figure 3F), which may help explain why Hsp104^R366E:D434R^ confers some induced thermotolerance in yeast, whereas Hsp104^R353E:E427K^ confers limited induced thermotolerance (Figure 3B).

Notably, Hsp104^R353E:E427R^ displayed WT levels of activity with Ssa1 and Sis1 (Figure 3E) and ∼42% Hsp104 activity with Ssa1 and Ydj1 (Figure 3F), which helps explain why this variant confers induced thermotolerance *in vivo* (Figure 3B), despite limited activity with human Hsc70 and DnaJA1 (Figure 3A). Indeed, it appears that the reconfigured NBD1:MD helix L1 salt bridge of Hsp104^R353E:E427K^, Hsp104^R353E:E427R^, and Hsp104^R366E:D434R^ creates a species barrier between yeast Hsp104 and human Hsc70 and DnaJA1, i.e., Hsp104^R353E:E427K^, Hsp104^R353E:E427R^, and Hsp104^R366E:D434R^ do not work well with human Hsc70 and DnaJA1 but are functional with yeast Ssa1 and Ydj1. Thus, it appears that the interprotomer interactions between NBD1 and MD helix L1 of the ATP state can be altered to establish species barriers between Hsp104 and Hsp70, which can occur naturally as with the barrier between yeast Hsp104 and bacterial Hsp70.^15,67^

Collectively, these results reveal the ATP-specific interprotomer interactions between NBD1 and MD helix L1 (Figure 1A) function as a rheostat to fine-tune Hsp104 collaboration with Hsp70 and Hsp40. Perturbation of these contacts can reduce Hsp70 and Hsp40 collaboration without potentiating activity (Figure 2A). Surprisingly, specific perturbation of this network yields hypomorphic Hsp104 variants that collaborate selectively with class B Hsp40s as with Hsp104^R419E^ and Hsp104^R366E^ (Figure 2A, C, D). Additionally, specific rewiring of this network potentiates Hsp104 activity as with Hsp104^E190K:R419E^ and Hsp104^E190R:R419E^ (Figure 3A). Rewiring the network in other ways can greatly reduce collaboration with Hsp70 and Hsp40 without affecting the intrinsic disaggregase activity of Hsp104 as with Hsp104^R194E:E412K^ and Hsp104^R366E:D434K^ (Figure 3C-F). Remarkably, reconfiguring the NBD1:MD helix L1 network in yet further ways can create species barriers with human Hsp70 as with Hsp104^R353E:E427R^ and Hsp104^R366E:D434R^ (Figure 3A, E, F). Hsp104^R353E:E427R^ and Hsp104^R366E:D434R^ operate more similarly to Hsp104 with the class B Hsp40, Sis1, but exhibit reduced activity with class A Hsp40, Ydj1 (Figure 3E, F), and confer up to ∼67% of WT Hsp104 levels of induced thermotolerance *in vivo* (Figure 3B). This finding suggests that collaboration with just Sis1 is insufficient for Hsp104 to confer induced thermotolerance *in vivo.* Indeed, collaboration with Sis1 and Ydj1 appears critical for WT levels of Hsp104 activity *in vivo.*^77^ Overall, we conclude that ATP-specific interprotomer interactions between NBD1 and MD helix L1 control several aspects of collaboration between Hsp104 and Hsp70 plus Hsp40.

### Perturbing the intraprotomer NBD1:MD contacts of the ADP state frequently potentiates Hsp104 activity

In contrast to the ATP state, the MD primarily interacts with NBD1 within the same subunit of the hexamer in the presence of ADP (Figure 1B). The role of these intraprotomer contacts in comparison to the interprotomer NBD1:MD contacts of the ATP state has remained unclear. Therefore, we explored 31 interactions involving 14 NBD1 residues and 19 MD residues in the interaction surface presented in Figure 1B. We designed 46 mutations aiming to alter these interactions, of which 16 have been tested previously.^29,30,39,58,59,63^ Thus, we generated 30 additional single missense Hsp104 variants expected to alter the interactions of this interface (Table S2 and Figure 1B).

We then assessed these Hsp104 variants for the ability to suppress the toxicity of α-synuclein, FUS, and TDP-43 in Δ*hsp104* yeast.^29^ Here, Hsp104 and the vector control are unable to mitigate α-synuclein, FUS, and TDP-43 toxicity, whereas positive control potentiated Hsp104 variants, Hsp104^A503V^or Hsp104^A503S^, strongly suppress toxicity.^29^ We find that ∼72% (33/46) of the Hsp104 variants designed to weaken the intraprotomer NBD1-MD interactions of the ADP state potentiate activity and enable mitigation of α-synuclein, FUS, and TDP-43 toxicity (Table S2). Among the 30 Hsp104 variants tested in this work, we found an additional 18 variants that enhanced activity and an additional 12 variants that did not (Figure 4 and Table S2). Among the 18 potentiated variants, only two variants (I361K and K470D) present off-target toxicity to yeast at 37°C (Figure S4A). Moreover, the expression level of Hsp104 variants in yeast is verified (Figure S4B) and all 30 Hsp104 variants are expressed. These results suggest that the intraprotomer NBD1-MD- interactions observed in the ADP state function to restrict Hsp104 activity. However, Hsp104 disaggregation activity can be unleashed by weakening these intraprotomer NBD1-MD- interactions, resulting in potentiated variants with low off-target toxicity.

**Figure 4.**
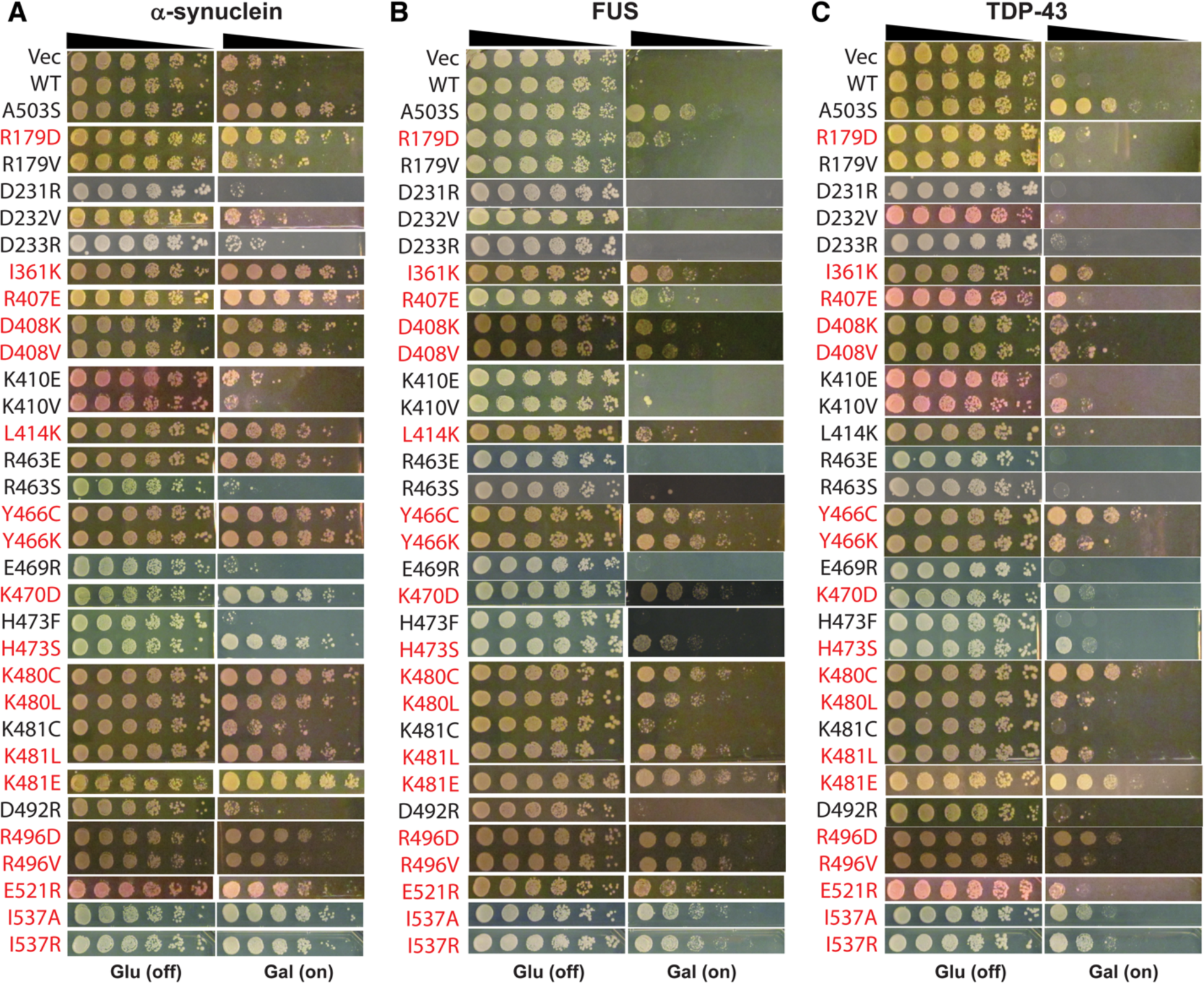
Perturbing the intraprotomer NBD1:MD contacts of the ADP state frequently potentiates Hsp104 activity. **(A-C)** Δ*hsp104* yeast integrated with α-synuclein-YFP (**A**) FUS (**B**), or TDP-43 (**C**) on a galactose-inducible promoter were transformed with the indicated Hsp104 variants that perturb the intraprotomer NBD1:MD contacts of the ADP state. Empty vector and WT Hsp104 are negative controls, and Hsp104^A503S^ is a positive control. Yeast were spotted onto glucose (induction is off) and galactose (induction is on) media in a five-fold serial dilution. Potentiated Hsp104 variants are highlighted in red. Western blots were performed to evaluate Hsp104 expression, see Figure S4. See also Figure S4 and Table S2.

Among the 18 potentiated variants discovered in this work, Hsp104^Y466C^, Hsp104^K480C^, Hsp104^K481E^, Hsp104^R496D^, Hsp104^I537A^, and Hsp104^I537R^ present strong activity in mitigating α- synuclein, FUS, and TDP-43 toxicity with minimal off-target toxicity, and resemble Hsp104^A503S^ (Figure 4 and S4A).^29^ By contrast, Hsp104^I361K^, Hsp104^Y466K^, Hsp104^K470D^, Hsp104^H473S^, Hsp104^K480L^, Hsp104^K481L^ and Hsp104^E521R^ show slightly reduced activity compared to Hsp104^A503S^ (Figure 4), whereas Hsp104^R179D^, Hsp104^R407E^, Hsp104^D408K^, and Hsp104^D408V^ were further reduced, but still can mitigate α-synuclein, FUS, and TDP-43 toxicity (Figure 4). Hsp104^L414K^ presents the lowest activity of the 18 new potentiated Hsp104 variants (Figure 4). Hsp104^L414K^ mitigates α-synuclein and FUS toxicity but is unable to reduce TDP-43 toxicity (Figure 4). Interestingly, a valine scan of the entire MD only identified Hsp104^R496V^ and Hsp104^K480V^ with potentiated activity among these same sites,^39^ highlighting the power of making rational, targeted mutations to potentiate activity.

### Restricting Hsp104 activity in the absence of Hsp70 reduces off-target toxicity of potentiated Hsp104 variants

Potentiated Hsp104 variants can exhibit unfavorable off-target toxicity, especially at 37°C in yeast.^29,30,63^ Hsp104 likely recognizes unfolded regions of proteins with a bias for peptides of a certain amino-acid composition rather than any specific sequence.^78^ Thus, it was proposed that potentiated Hsp104 variants may recognize and unfold metastable proteins, which results in toxicity.^29,63^ One concept is that Hsp70 may direct Hsp104 to aggregated structures and away from soluble misfolded polypeptides or naturally metastable proteins.^66^ Since potentiated Hsp104 variants can function independently of Hsp70 to varying extents,^29,30,63,64^ this mechanism of substrate selection may become dysregulated such that excessive soluble polypeptide unfolding drives off-target toxicity. However, WT Hsp104 is too tightly regulated and is unable to overcome widespread aggregation by neurodegenerative disease proteins such as TDP-43, FUS, and α- synuclein in yeast.^29,30,63,64^ Thus, we hypothesize that potentiated Hsp104 variants with reduced unfoldase activity for soluble proteins, and partial independence from Hsp70 may reside in an advantageous therapeutic window. These Hsp104 variants would preferentially target aggregated proteins without excessive and toxic unfolding of soluble proteins. Thus, we set out to test whether potentiated Hsp104 variants with greater off-target toxicity have stronger unfoldase activity against soluble proteins and less dependence on Hsp70 for protein disaggregation.

To test this concept, we selected several Hsp104 variants with a range of off-target toxicities: Hsp104 (no off-target toxicity), Hsp104^S535E^ (an MD variant with minimal off-target toxicity), Hsp104^E360R^ (an NBD1 variant with minimal off-target toxicity), and Hsp104^I187F^ (an NBD1 variant with significant off-target toxicity) (Figure 5A).^29,63,64^ We then assessed the ability of these Hsp104 variants to unfold the model substrate RepA_1-15_-GFP, which is comprised of the N-terminal 15 residues of RepA appended to the N-terminus of GFP.^79^ The RepA_1-15_ tag is a short, unfolded region, which is sufficient to target RepA_1-15_-GFP for unfolding by potentiated Hsp104 variants. By contrast, Hsp104 does not unfold RepA_1-15_-GFP in the presence of ATP (Figure 5B). Strikingly, however, we find Hsp104^I187F^, which has the most off-target toxicity, unfolds RepA_1-15_-GFP the most rapidly, whereas the less toxic Hsp104 variants, Hsp104^S535E^ and Hsp104^E360R^ unfold RepA_1- 15_-GFP less rapidly than Hsp104^I187F^ (Figure 5B). Hence, off-target toxicity correlates positively with stronger unfoldase activity against soluble protein.

**Figure 5.**
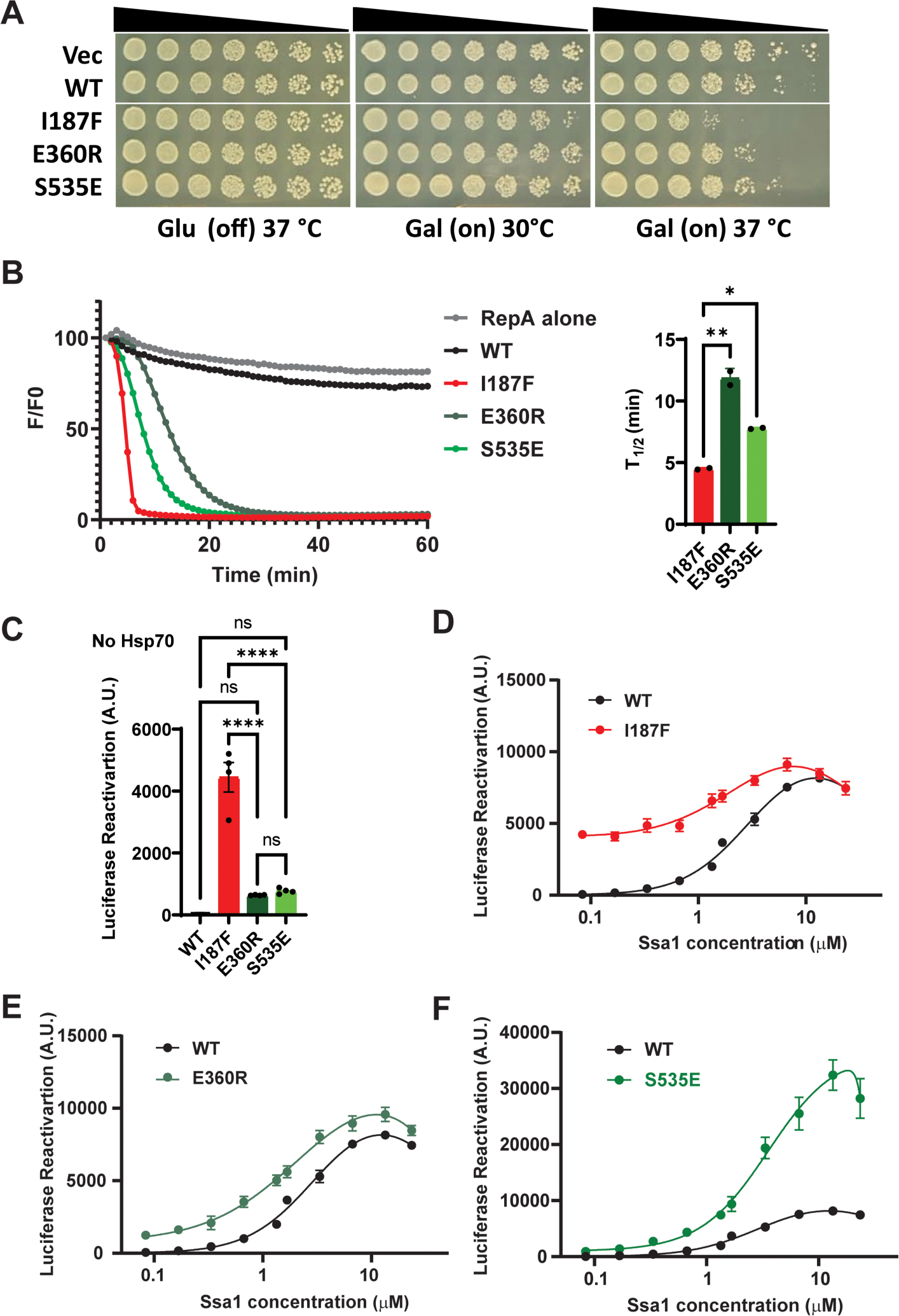
Restricting Hsp104 activity in the absence of Hsp70 reduces off-target toxicity of potentiated Hsp104 variants. **(A)** *Δhsp104* yeast were transformed with the indicated galactose-inducible Hsp104 variants or an empty vector or Hsp104 control. Yeast were spotted onto glucose (induction off) and galactose (induction on) media in a five-fold serial dilution. Yeast were incubated on a glucose plate 37°C (left) or on a galactose plate at 30°C (middle) or 37°C (right). Note that Hsp104^I187F^ is more toxic than Hsp104^E360R^ or Hsp104^S535E^. Hsp104^E360R^ and Hsp104^S535E^ are similar to Hsp104 at 37°C on galactose. **(B)** RepA_1-15_-GFP (0.7µM) unfolding activity of the indicated Hsp104 variant (6µM, monomeric) in the presence of ATP (4mM) and GroEL_trap_ (2.5µM). RepA_1-15_-GFP unfolding (%) was assessed by the RepA_1-15_-GFP fluorescence signal at the indicated time (F), divided by the RepA_1-15_-GFP fluorescence signal at time 0 (F0). Left, kinetics of RepA_1-15_-GFP unfolding. Results from a representative experiment are shown. Right, the half time of RepA_1-15_-GFP unfolding for each Hsp104 variant. Bars represent means±SEM (N=2), each replicate is shown as a dot. One-way ANOVA Tukey test was performed to compare the half-time of one Hsp104 variant to every other one at 95% CI. *P=0.0117, **P=0.0012 for I187F vs. E360R, **P=0.0063 for E360R vs. S535E. (**C**) Luciferase disaggregation and reactivation by Hsp104, Hsp104^I187F^, Hsp104^E360R^, or Hsp104^S535E^ in the absence of Hsp70 and Hsp40. The indicated Hsp104 variant (1µM, monomeric) was incubated with chemically denatured luciferase aggregates (100 nM monomer concentration) for 90 min. Bars represent means±SEM (N=4), each replicate is shown as a dot. One-way ANOVA Tukey test was performed to compare the level of reactivated luciferase aggregates by Hsp104 variants at 95% CI. **** P≤0.0001. **(D-F)** Luciferase disaggregation and reactivation by Hsp104 (black curve), Hsp104^I187F^ (D, red curve), Hsp104^E360R^ (E, green curve) and Hsp104^S535E^ (F, green curve) as a function of Ssa1 concentration as indicated on the x-axis (log scale). The indicated Hsp104 variant (1µM, monomeric) in the presence of various Ssa1 concentrations was incubated with chemically denatured luciferase aggregates (100 nM monomer concentration). Values represent means±SEM (N=2). A bell-shaped dose-dependent curve is used to fit the data; see STAR Methods.

To measure the Hsp70 dependence for Hsp104 disaggregase activity, we performed luciferase disaggregation and reactivation experiments in the absence of Hsp70 or Hsp40 (Figure 5C). Here, we find that Hsp104^I187F^ has the highest luciferase disaggregation and reactivation activity in the absence of Hsp70 or Hsp40 (Figure 5C). By contrast, Hsp104^E360R^ and Hsp104^S535E^ show significantly lower activity in the absence of Hsp70 and Hsp40, whereas Hsp104 is completely inactive as expected (Figure 5C).^15^ Thus, off-target toxicity correlates positively with stronger disaggregase activity in the absence of Hsp70 or Hsp40.

Next, we titrated Ssa1 (in the absence of Hsp40) into luciferase disaggregation and reactivation experiments with each potentiated Hsp104 variant versus Hsp104. Hsp104^I187F^ shows very little dependence on Ssa1 compared to Hsp104 (Figure 5D). By contrast, Hsp104^E360R^ shows stronger dependence on Ssa1, but outperforms Hsp104 at every Hsp70 concentration tested (Figure 5E). Intriguingly, Hsp104^S535E^ also shows Ssa1 dependence, but is greatly stimulated at high Ssa1 concentrations (Figure 5F). Thus, potentiated Hsp104 variants with minimal off-target toxicity are also more dependent on Hsp70 in protein disaggregation.

### Rationally designed potentiated Hsp104 with minimized off-target toxicity

Next, we developed Hsp104 variants with minimized off-target toxicity based on the design principles established above, i.e., reduced unfoldase activity for soluble proteins and partial independence from Hsp70 for protein disaggregation. Given that the ATP-specific interprotomer NBD1:MD helix L1 interface functions as a rheostat for collaboration with Hsp70 and Hsp40 (Figure 2, 3), we focused on this region to define modifications that tune Hsp70-Hsp40 collaboration to the desired level.

Upon closer inspection of this salt bridge interaction, we observed that E190, E191, and E192 of NBD1 are positioned such that they could all potentially contact R419 (Figure 6A, left panel). E190, E191, and E192 are located at a loop between Helix B2 and B3 in NBD1 that forms a junction with the coiled-coil MD helix L1 (Figure 6A, left panel). This loop may provide enough freedom for these residues to have a dynamic interaction with R419. To test if salt-bridge interactions between E190, E191, or E192 and R419 can regulate Hsp104 activity and collaboration with Hsp70 and Hsp40, we designed variants to rewire these interactions and modify MD orientation. Thus, we explored combinations of R419E with arginine substitutions at E190, E191, or E192. We tested whether these designed variants could mitigate α-synuclein, FUS, and TDP-43 toxicity in yeast and whether they exhibited off-target toxicity. All the Hsp104 variants were expressed at roughly similar levels and did not affect disease protein expression (Figure S5).

**Figure 6.**
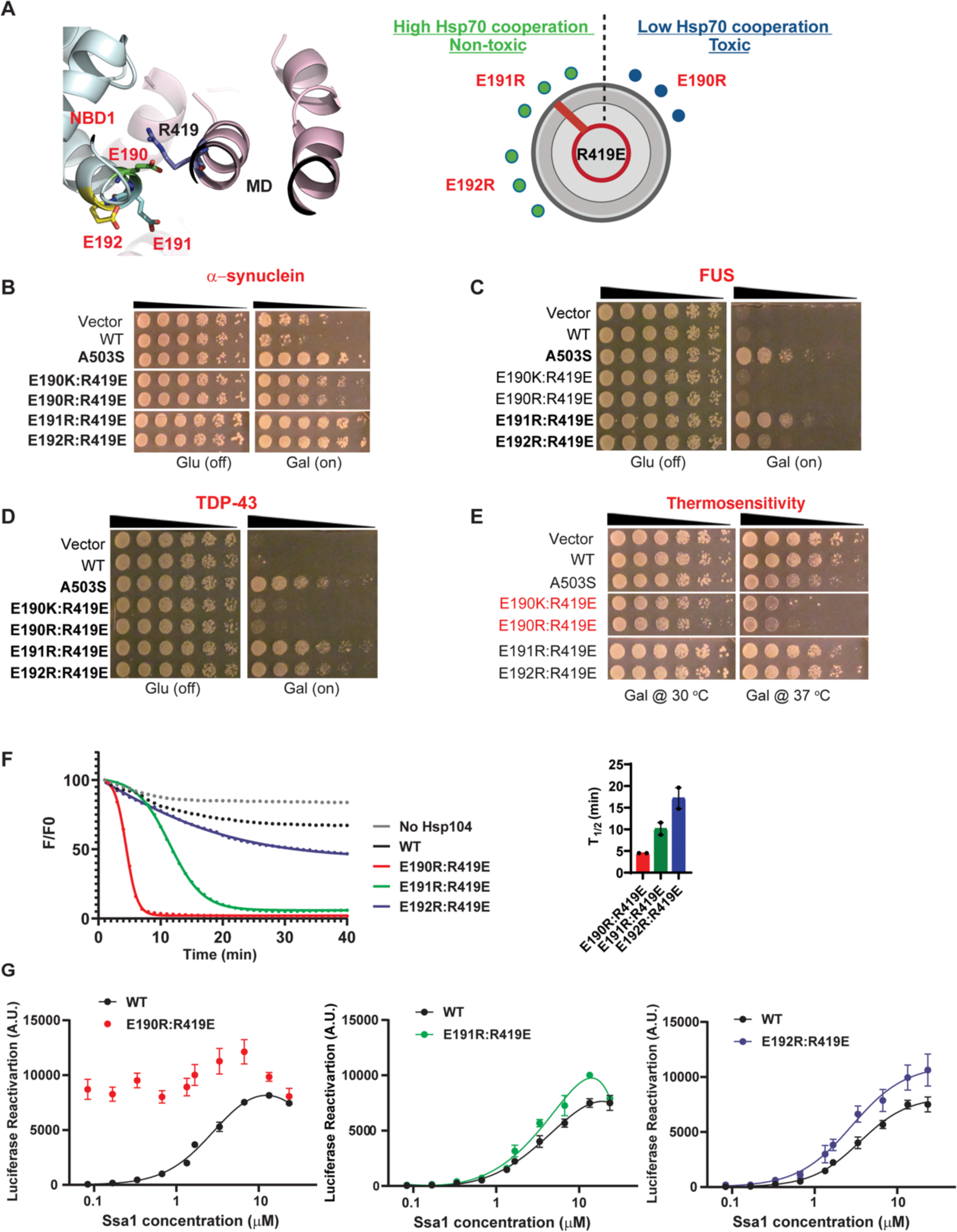
Rational design of potentiated Hsp104 variants with minimized off-target toxicity. (**A**) NBD1 (blue) residues E190, E191, and E192 (in red) form a rheostat-like interaction with R419 (black) in the MD (colored in pink) of the adjacent subunit. Hsp104 variants are designed to alter these interactions like a rheostat (right panel) to tune Hsp104- Hsp70 interaction to a suitable level of potentiated activity without off-target toxicity. **(B-D)** Spotting assay testing the ability of Hsp104 variants to mitigating α-synuclein (B), FUS (C), and TDP-43 (D) toxicity in yeast. *Δhsp104* yeast integrated with α-synuclein (B), FUS (C), and TDP- 43 (D) on a galactose-inducible promoter were transformed with Hsp104 variants or an empty vector, WT Hsp104 or Hsp104^A503S^ controls. Yeast were spotted onto glucose (induction off) and galactose (induction on) media in a five-fold serial dilution. The variants that have potentiated activity to mitigate disease protein toxicity in yeast are highlighted in bold. (**E**) The toxicity of designed Hsp104 variants in yeast were evaluated at 37°C. *Δhsp104* yeast were transformed with galactose-inducible Hsp104 variants or an empty vector, WT Hsp104 or Hsp104^A503S^ controls. The yeast were spotted onto glucose (induction off) and galactose (induction on) media in a five-fold serial dilution. Yeast were incubated at 30°C (left) or 37°C (right) on galactose plates. The toxic variants are highlighted in red. (**F**) RepA_1-15_-GFP (0.7µM) unfolding kinetics by Hsp104^E190R:R419E^, Hsp104^E191R:R419E^, or Hsp104^E192R:R419E^ (6μM, monomeric concentration) is measured in the presence of a GroEL_trap_ (2.5μM). RepA_1-15_-GFP unfolding (%) was assessed by the RepA_1-15_-GFP fluorescence signal at the indicated time (F), divided by the RepA_1-15_-GFP fluorescence signal at time 0 (F0). Results from a representative experiment are shown. The half-time of RepA_1-15_-GFP unfolding for each Hsp104 is shown on the right. Bars represent means±SEM (N=2), each replicate is shown as a dot. (**G**) Luciferase disaggregation and reactivation by Hsp104 (black curve), Hsp104^E190R:R419E^ (red curve, left), Hsp104^E191R:R419E^ (red curve, middle) and Hsp104^E192R:R419E^ (red curve, right) as a function of Ssa1 concentration. The indicated Hsp104 variant (1µM, monomeric) in the presence of various Ssa1 concentrations as indicated on the x-axis (log scale) was incubated with chemically denatured luciferase aggregates (100nM monomer concentration) for 90 min. Values represent means±SEM (N=2). A bell-shaped dose-dependent curve is used to fit the data, see STAR Methods. See also Figure S5.

Previously, we found that Hsp104^E190K:R419E^ and Hsp104^E190R:R419E^ have enhanced disaggregase activity (Figure 3A). However, this activity was largely independent of Hsp70 and Hsp40, which predicts off-target toxicity (Figure 3A). In yeast, we found that these Hsp104 variants weakly suppressed α-synuclein toxicity (Figure 6B) but were unable to suppress FUS toxicity (Figure 6C), and very weakly suppressed TDP-43 toxicity (Figure 6D). This pattern of disease protein toxicity mitigation is unusual and has not been observed before.^29,30,37,58,63,64^ Moreover, as predicted, these variants exhibited off-target toxicity at 37°C (Figure 6E). Hence, the E190K/R:R419E variants have the rheostat dialed too far toward Hsp70 independence (Figure 6A, right panel).

By contrast, Hsp104^E191R:R419E^ and Hsp104^E192R:R419E^ strongly suppress α-synuclein, FUS, and TDP-43 toxicity (Figure 6B-D). Importantly, these variants did not exhibit off-target toxicity (Figure 6E). Hence, the E191R:R419E and E192R:R419E variants have the rheostat dialed to an appropriate level of Hsp70 collaboration (Figure 6A, right panel).

Based upon these observations, we would predict that Hsp104^E190R:R419E^ would unfold RepA_1-15_- GFP more rapidly than Hsp104^E191R:R419E^ and Hsp104^E192R:R419E^. We would also predict that Hsp104^E190R:R419E^ would be less dependent on Hsp70 than Hsp104^E191R:R419E^ and Hsp104^E192R:R419E^ for luciferase disaggregation and reactivation. These predictions were confirmed experimentally (Figure 6F, G). Collectively, these findings suggest that interactions between NBD1 residues E190, E191, or E192 and MD helix L1 R419 function as a rheostat that can be adjusted to fine-tune collaboration with Hsp70.

### Potentiated Hsp104 variants suppress FUS proteinopathy in human cells

A feature of degenerating neurons in various FUS proteinopathies, including ALS and frontotemporal dementia (FTD), is the depletion of FUS from the nucleus and the accumulation of FUS in cytoplasmic inclusions.^10^ Potentiated Hsp104 variants can mitigate cytoplasmic FUS aggregation and toxicity in yeast,^29^ but their activity in human cells has not been assessed. To this end, we utilized a human (HeLa) cell model by expressing mCherry-tagged Hsp104 and GFP-tagged FUS in HeLa cells.^80^ We selected a panel of potentiated variants with minimal off-target toxicity to assess in this system. Importantly, potentiated Hsp104 variants can significantly suppress FUS cytoplasmic mislocalization and aggregation, whereas Hsp104 cannot (Figure 7A). Indeed, Hsp104^A503S^, Hsp104^E191R:R419E^, Hsp104^E360R^, Hsp104^K481E^, and Hsp104^S535E^ suppressed cytoplasmic FUS aggregation and maintained FUS in the nucleus (Figure 7A, B) without reducing FUS expression level (Figure 7C). Thus, enhanced Hsp104 variants provide a mechanism to mitigate aberrant cytoplasmic FUS aggregation in human cells.

**Figure 7.**
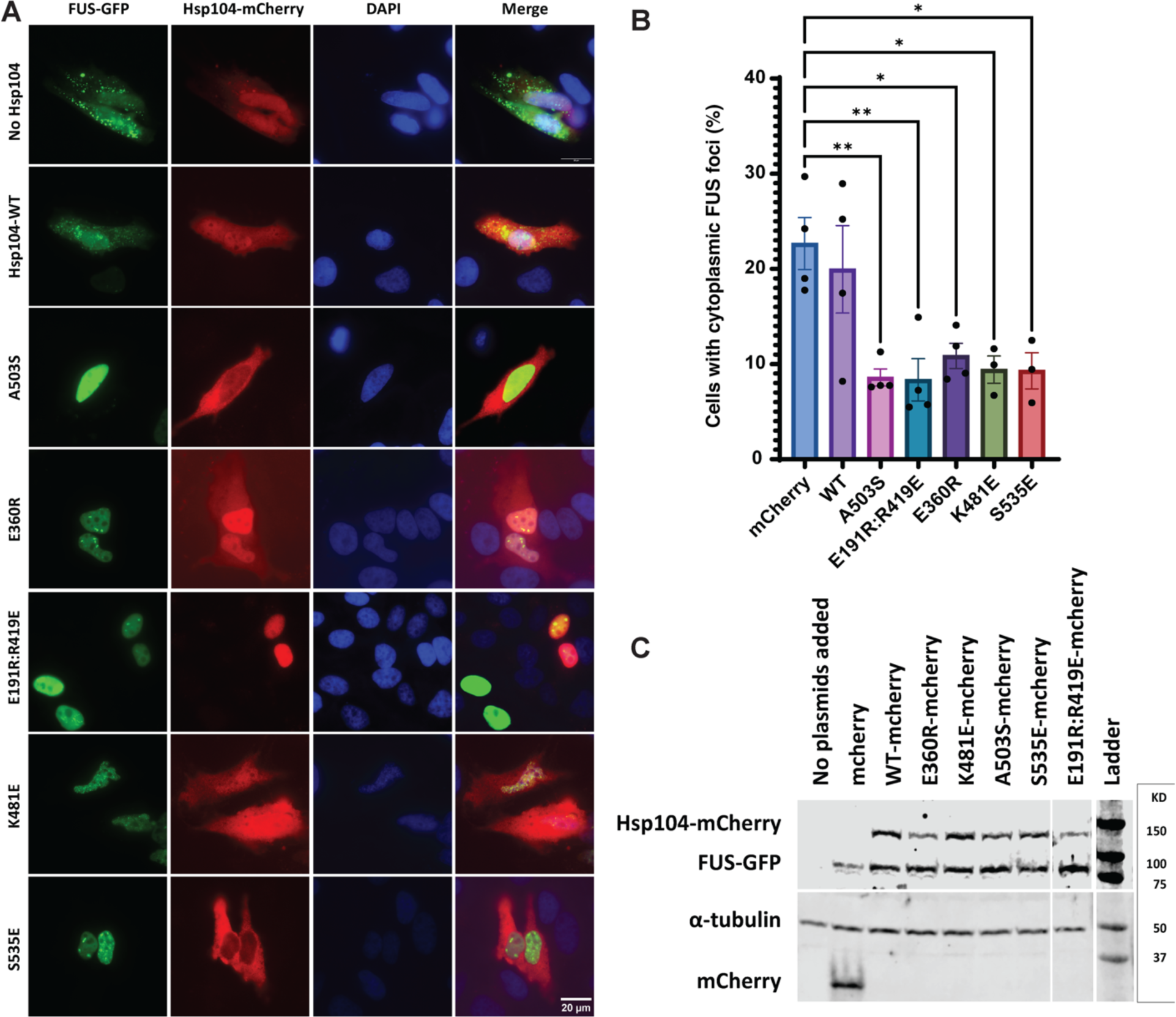
Potentiated Hsp104 variants suppress FUS proteinopathy in human cells. (**A**) After 24h, HeLa cells transfected with GFP-FUS (Green) and mCherry, or the indicated Hsp104 variant-mCherry (Red) were fixed and imaged. DAPI is used to stain the nucleus. Representative images are shown. Scale bar, 20μm. (**B**) Quantification of the percentage of cells with cytoplasmic FUS foci. 500 to 800 cells were counted over four separate trials for each Hsp104 variant and mCherry control; data points represent independent transfections. Bars represent means±SEM (N=4). One-way ANOVA with correction for multiple comparisons by Dunnett’s test was performed at 95% CI. **P≤0.01; *P≤0.05. (**C**) Western blot of lysates of HeLa cells transfected with GFP-FUS and either mCherry alone or mCherry-tagged Hsp104. Probing with anti-GFP and anti-mCherry shows FUS-GFP and Hsp104-mCherry expression levels, respectively. α-tubulin is used as a loading control.

## Discussion

In this work, we performed an intensive structure-function analysis of how ATP-specific or ADP-specific MD configurations regulate Hsp104 disaggregase activity. We determined that the ATP-specific interactions between MD helix L1 and NBD1 of the adjacent clockwise protomer are critical for Hsp104 to collaborate effectively with Hsp70 and Hsp40 during protein disaggregation. Specifically, salt-bridge interactions between NBD1:MD L1 via E190:R419, R194:E412, R353:E427, and R366:D434, enable Hsp104 to collaborate with Hsp70 and Hsp40. Intriguingly, disrupting these interactions does not potentiate activity or affect the intrinsic disaggregase activity of Hsp104. Thus, Hsp104 can still couple ATP hydrolysis to substrate processing when these contacts are broken. However, the ability of Hsp104 to collaborate with Hsp70 and Hsp40 is specifically disrupted. These findings are surprising as it was anticipated that these interactions would be important for intersubunit collaboration within the hexamer rather than collaboration with Hsp70 and Hsp40.

Intriguingly, we find that specific perturbations of the ATP-specific NBD1:MD helix L1 interactions (i.e., R419E or R366E) yielded hypomorphic Hsp104 variants. These Hsp104 hypomorphs confer some thermotolerance *in vivo*. In luciferase disaggregation and reactivation *in vitro*, Hsp104^R419E^ and Hsp104^R366E^ work selectively with Ssa1 and Sis1, and are unable to function with Ssa1 and Ydj1 or human Hsc70 and DnaJA1. Thus, these Hsp104 variants displayed selectivity to function with the class B Hsp40, Sis1, but not class A Hsp40s, Ydj1 or DnaJA1. Hsp104^R419E^ and Hsp104^R366E^ were also less able to directly collaborate with Ssa1 directly (i.e., in the absence of Hsp40) and were inhibited by Ydj1. These findings emphasize the importance of ATP-specific NBD1:MD helix L1 interactions for productive collaboration with Hsp70 and class A Hsp40s. Moreover, they reinforce the importance of Hsp104 collaboration with Ssa1 and Ydj1 for thermotolerance *in vivo.*^72,75,76^

Having identified the critical importance of ATP-specific NBD1:MD helix L1 interprotomer interactions, we next assessed the consequences of rewiring these connections. Notably, rebuilding the E190:R419 salt bridge to E190R:R419E or E190K:R419E yielded potentiated Hsp104 variants, which functioned independently of Hsp70. Remarkably, reconfiguring the NBD:MD helix L1 network in yet further ways can create species barriers with human Hsp70 as with Hsp104^R353E:E427R^ and Hsp104^R366E:D434R^. Thus, manipulating the network of ATP-specific NBD1:MD helix L1 interprotomer interactions can: (a) reduce Hsp70 collaboration without enhancing activity; (b) generate hypomorphic Hsp104 variants that collaborate selectively with class B Hsp40s; (c) produce Hsp70-independent potentiated variants; or (d) create species barriers between Hsp104 and Hsp70. Collectively, these findings suggest that the ATP-specific NBD1: MD helix L1 interactions function as a rheostat to tune the level of collaboration with Hsp70 and Hsp40. Indeed, the ATP-specific network of interprotomer contacts between NBD1 and MD helix L1 appears to be poised as a capacitor that can release diverse phenotypes.

By contrast, the ADP-specific intraprotomer contacts between NBD1 and MD helix L2 function to restrict activity. When these contacts were disrupted (Figure 4 and Table S2), we observed enhanced Hsp104 activity in mitigating α-syn, FUS and TDP-43 toxicity in yeast, indicating enhanced disaggregase activity of these variants. Indeed, we designed 46 variants to alter 31 ADP-specific intraprotomer contacts, and 33 variants exhibited potentiated activity (Figure 4 and Table S2). Disrupting these contacts likely increases the rate of ADP release from NBD1, which accelerates the Hsp104 motor.^40^ Notably, many residues, such as D233, E360, E366, E412, and R419, are involved in both intraprotomer and interprotomer contacts between NBD1 and the MD, indicating a dynamic and highly regulated network of interactions, which likely enable communication within and between subunits during disaggregation.

A difficulty in developing potentiated Hsp104 variants as therapeutic agents lies in their off-target toxicity, which likely stems from their ability to unfold, metastable soluble proteins or soluble proteins with partially unfolded regions.^36^ One solution to this problem is to increase the substrate specificity of potentiated Hsp104 variants for specific neurodegenerative-disease proteins, which we have achieved with α-synuclein.^30^ However, multiple proteins can aggregate in neurodegenerative disease, which may limit the utility of substrate-specific protein disaggregases. Another strategy would be to tune Hsp104 activity such that potentiated disaggregase activity is retained while unfolding of soluble proteins is minimized. Here, we establish that fine-tuning the level of Hsp70 collaboration provides a mechanism to achieve this goal. We reach this conclusion by first considering three potentiated Hsp104 variants: Hsp104^I187F^, Hsp104^E360R^, and Hsp104^S535E^.^63,64^ Hsp104^I187F^ exhibits more off-target toxicity than Hsp104^E360R^, which in turn exhibits more off-target toxicity than Hsp104^S535E^.^63,64^ Strikingly, Hsp104^I187F^ unfolds RepA_1-15_- GFP (a model soluble protein with a partially unfolded region) more rapidly than Hsp104^E360R^, which in turn unfolds RepA_1-15_-GFP more rapidly than Hsp104^S535E^. Furthermore, Hsp104^I187F^ displays less dependence on Hsp70 than Hsp104^E360R^ or Hsp104^S535E^ in luciferase disaggregation and reactivation. Thus, too much independence from Hsp70 may yield off-target toxicity, whereas too much dependence of Hsp70 (as with wild-type Hsp104) leads to a reduced ability to combat deleterious protein misfolding connected with neurodegenerative disease. Overall, our findings suggest rules for minimizing off-target toxicity: (1) minimize the ability of the Hsp104 variant to unfold soluble proteins with partially unfolded regions; and (2) tune the level of collaboration with Hsp70.

We then leveraged this knowledge to adjust the ATP-specific NBD1:MD helix L1 rheostat to the appropriate level of Hsp70 collaboration. We find that Hsp104^E190R:R419E^ displays increased off-target toxicity, enhanced ability to unfold RepA_1-15_-GFP, and less dependence on Hsp70 in luciferase disaggregation and reactivation. By contrast, Hsp104^E191R:R419E^ and Hsp104^E192R:R419E^ display reduced off-target toxicity, reduced ability to unfold RepA_1-15_-GFP, and more dependence on Hsp70 in luciferase disaggregation and reactivation. Hence, these NBD1:MD helix L1 variants may set the ATP-specific MD configuration in a way that optimally tunes Hsp70 collaboration to yield potentiated Hsp104 variants with minimal off-target toxicity.

Finally, we establish that potentiated Hsp104 variants with minimal off-target effects can mitigate aberrant FUS aggregation in human cells for the first time. Thus, a panel of potentiated Hsp104 variants can reduce cytoplasmic FUS aggregation in human cells, whereas Hsp104 is ineffective. These findings suggest that Hsp104 and enhanced variants can be translated to reduce deleterious protein aggregation in human cells, which sets the stage for further developing Hsp104 as a therapeutic agent. Indeed, in this light, advances in lipid nanoparticle-mediated mRNA therapeutics are particularly exciting as they provide a mechanism to introduce a transient dose of Hsp104 variants to where they are needed.^81,82^ In this way, potentiated Hsp104 variants could relieve toxic protein aggregation and then be downregulated such that potential off-target effects are minimized.

## STAR Methods

### Key resources table

| REAGENT or RESOURCE | SOURCE | IDENTIFIER |
| --- | --- | --- |
| **Antibodies** |  |  |
| Rabbit anti-Hsp104 polyclonal | Enzo Life Sciences | Cat# ADI-SPA-1040-F; RRID: AB_2039208 |
| Rabbit anti-FUS polyclonal | Bethyl Laboratories | Cat# A300-302A; RRID: AB_309445 |
| Rabbit anti-TDP-43 polyclonal | Proteintech | Cat# 10782; RRID: AB_615042 |
| Rabbit anti-GFP polyclonal | Sigma-Aldrich | Cat# G1544; RRID: AB_439690 |
| Rabbit monoclonal anti-alpha tubulin antibody | Abcam | Cat# ab6160 |
| Mouse anti-PGK1 monoclonal | Thermo Fisher | Cat# 459250; RRID: AB_2532235 |
| Anti-mCherry polyclonal antibody | Abcam | Cat# ab167453 |
| IRDye 800CW Goat anti-Mouse IgG secondary antibody | LI-COR | Cat# 926-32210; RRID:AB_621842 |
| IRDye 680RD Goat anti-Rabbit IgG secondary antibody | LI-COR | Cat# 926-68071; RRID: AB_10956166 |
| IRDye 800CW Goat anti-Rat IgG Secondary Antibody | LI-COR | Cat#926-32219; RRID: AB_1850025 |
| **Bacterial and virus strains** |  |  |
| *Escherichia coli* DH5α competent cells | Thermo Fisher | Cat# 18265017 |
| *Escherichia coli* BL21-CodonPlus (DE3) -RIL competent cells | Agilent | Cat# 230245 |
| One Shot TOP10 Chemically Competent *Escherichia coli* | Invitrogen | Cat# C404010 |
| **Chemicals, peptides, and recombinant proteins** |  |  |
| Creatine phosphate | Roche | Cat# 10621722001 |
| Adenosine 5'-triphosphate disodium salt hydrate (ATP) | Sigma-Aldrich | Cat# A3377 |
| Adenosine 5'-[γ-thio] triphosphate tetralithium salt | Roche | Cat# 11162306001 |
| DpnI | NEB | Cat# R0176S |
| Gateway BP Clonase II Enzyme Mix | Invitrogen | Cat# 11789013 |
| Gateway LR Clonase II Enzyme Mix | Invitrogen | Cat# 11791019 |
| NEBuilder HiFi DNA Assembly Master Mix | NEB | Cat# E2621S |
| cOmplete, Mini, EDTA-free Protease Inhibitor Cocktail | Sigma-Aldrich | Cat# 4693159001 |
| Casein fluorescein isothiocyanate from bovine milk (FITC-Casein) | Sigma-Aldrich | Cat# C0528 |
| Creatine kinase | Roche | Cat# 10127566001 |
| Firefly luciferase | Sigma-Aldrich | Cat# L9506 |
| Lysozyme | Sigma-Aldrich | Cat# L6876 |
| Hoechst 33342 stain | Invitrogen | Cat# H3570 |
| Lipofectamine 2000 Transfection Reagent | Invitrogen | Cat# 11668027 |
| VECTASHIELD Antifade Mounting Medium with DAPI | Vector Laboratories | Cat# H-1200-10 |
| DMEM, high glucose, pyruvate | Gibco | Cat# 11995065 |
| Fetal bovine serum | Cytiva | Cat# SH30910.03 |
| Penicillin-streptomycin | Gibco | Cat# 15140122 |
| TeSR Plus medium | STEMCELL Technologies | Cat# 100-0276 |
| HCl | Sigma | Cat# H9892 |
| His-TEV protease | Cupo and Shorter<sup>83</sup> | N/A |
| Hsp104 | Jackrel et al.<sup>29</sup> | N/A |
| Hsp104<sup>A503S</sup> | Jackrel et al.<sup>29</sup> | N/A |
| Hsp104<sup>R179D/V</sup> | This paper | N/A |
| Hsp104<sup>I187F</sup> | Tariq et al.<sup>63</sup> | N/A |
| Hsp104<sup>E190R/K</sup> | This paper | N/A |
| Hsp104<sup>E190R/K;R419E</sup> | This paper | N/A |
| Hsp104<sup>E191R;R419E</sup> | This paper | N/A |
| Hsp104<sup>E192R:R419E</sup> | This paper | N/A |
| Hsp104<sup>E190R:E191R:R419E</sup> | This paper | N/A |
| Hsp104<sup>E190R:E192R:R419E</sup> | This paper | N/A |
| Hsp104<sup>R194E</sup> | Gates et al.<sup>59</sup> | N/A |
| Hsp104<sup>E412K</sup> | Gates et al.<sup>59</sup> | N/A |
| Hsp104<sup>R194E:E412K</sup> | This paper | N/A |
| Hsp104<sup>R353E</sup> | Gates et al.<sup>59</sup> | N/A |
| Hsp104<sup>E427K</sup> | Gates et al.<sup>59</sup> | N/A |
| Hsp104<sup>R353E:E427K</sup> | This paper | N/A |
| Hsp104<sup>R366E</sup> | Gates et al.<sup>59</sup> | N/A |
| Hsp104<sup>D434K</sup> | Gates et al.<sup>59</sup> | N/A |
| Hsp104<sup>R366E:D434K</sup> | This paper | N/A |
| Hsp104<sup>D231R</sup> | This paper | N/A |
| Hsp104<sup>D232V/R</sup> | This paper | N/A |
| Hsp104<sup>D233R</sup> | This paper | N/A |
| Hsp104<sup>K358D</sup> | Mack et al.<sup>30</sup> | N/A |
| Hsp104<sup>E360R</sup> | Tariq et al.<sup>63</sup> | N/A |
| Hsp104<sup>I361K</sup> | This paper | N/A |
| Hsp104<sup>D408K/V</sup> | This paper | N/A |
| Hsp104<sup>K410E/L/V</sup> | This paper | N/A |
| Hsp104<sup>D492R</sup> | This paper | N/A |
| Hsp104<sup>R495D/E/N/M</sup> | Tariq et al.<sup>63</sup> | N/A |
| Hsp104<sup>R496D</sup> | This paper | N/A |
| Hsp104<sup>R496V</sup> | Ryan et al.<sup>39</sup> | N/A |
| Hsp104<sup>D484K</sup> | Mack et al.<sup>30</sup> | N/A |
| Hsp104<sup>K480E</sup> | Tariq et al.<sup>63</sup> | N/A |
| Hsp104<sup>K480C/L</sup> | This paper | N/A |
| Hsp104<sup>K481C/E/L</sup> | This paper | N/A |
| Hsp104<sup>Y466C/K</sup> | This paper | N/A |
| Hsp104<sup>E521R/V</sup> | This paper | N/A |
| Hsp104<sup>M536K</sup> | This paper | N/A |
| Hsp104<sup>I537A/R</sup> | This paper | N/A |
| Hsp104<sup>H473F</sup> | This paper | N/A |
| Hsp104<sup>A531V</sup> | This paper | N/A |
| Hsp104<sup>R407E</sup> | This paper | N/A |
| Hsp104<sup>K470D/V</sup> | This paper | N/A |
| Hsp104<sup>K470Q</sup> | Jackrel et al.<sup>58</sup> | N/A |
| Hsp104<sup>D408K/V</sup> | This paper | N/A |
| Hsp104<sup>L414K</sup> | This paper | N/A |
| Hsp104<sup>N539L/E/D/G/K</sup> | Jackrel et al.<sup>29</sup> | N/A |
| Hsp104<sup>P411H</sup> | This paper | N/A |
| Hsp104<sup>E412K</sup> | This paper | N/A |
| Hsp104<sup>R419E</sup> | Gates et al.<sup>59</sup> | N/A |
| GroEL<sub>trap</sub> | Jackrel et al.<sup>29</sup> | N/A |
| RepA<sub>1-15</sub>-GFP | This paper | N/A |
| Hsc70 | Enzo Life Sciences | Cat#ADI-SPP-751-F |
| DnaJA1 | Enzo Life Sciences | Cat#ADI-SPP-405-F |
| pE-His<sub>6</sub>-SUMO-Ssa1 | Michalska et al.<sup>84</sup> | N/A |
| pE-His<sub>6</sub>-SUMO-Sis1 | Michalska et al.<sup>84</sup> | N/A |
| pE-His<sub>6</sub>-SUMO-Ydj1 | Michalska et al.<sup>84</sup> | N/A |
| pFGET19-Ulp1 | Addgene | Plasmid #64697 |
| Luciferase Assay Reagent | Promega | Cat# E1483 |
| ATPase Activity Kit (Colorimetric) | Innova Biosciences | Cat# 601–0120 |
| QuikChange Site-Directed Mutagenesis Kit | Agilent | Cat# 200518 |
| **Experimental models: Organisms/strains** |  |  |
| *S. cerevisiae*: W303aΔ*hsp104* (*MATa*, *can1-100*, *his3-11, 15*, *leu2-3, 112*, *trp1-1*, *ura3-1*, *ade2-1*, *hsp104::KanMX*) | Jackrel et al.<sup>29</sup> | N/A |
| *S. cerevisiae: W303aΔhsp104-pAG303GAL-α-syn-YFP-pAG304GAL-α-syn-YFP* | Jackrel et al.<sup>29</sup> | N/A |
| *S. cerevisiae: W303aΔhsp104-pAG303GAL-FUS* | Jackrel et al.<sup>29</sup> | N/A |
| *S. cerevisiae: W303aΔhsp104-pAG303GAL-TDP-43* | Jackrel et al.<sup>29</sup> | N/A |
| *Cell line:* |  |  |
| Human:HeLa | ATCC | Cat#CCL-2;RRID:CVCL_0030 |
| **Recombinant DNA** |  |  |
| pAG416GAL-CCDB |  |  |
| pAG416GAL-Hsp104 | Jackrel et al.<sup>29</sup> | N/A |
| pAG416GAL-Hsp104<sup>A503S</sup> | Jackrel et al.<sup>29</sup> | N/A |
| pAG416GAL-Hsp104<sup>R179D/V</sup> | This paper | N/A |
| pAG416GAL-Hsp104<sup>I187F</sup> | Tariq et al.<sup>63</sup> | N/A |
| pAG416GAL-Hsp104<sup>E190R/K</sup> | This paper | N/A |
| pAG416GAL-Hsp104<sup>E190R/K:R419E</sup> | This paper | N/A |
| pAG416GAL-Hsp104<sup>E191R:R419E</sup> | This paper | N/A |
| pAG416GAL-Hsp104<sup>E192R:R419E</sup> | This paper | N/A |
| pAG416GAL-Hsp104<sup>E190R:E191R:R419E</sup> | This paper | N/A |
| pAG416GAL-Hsp104<sup>E190R:E192R:R419E</sup> | This paper | N/A |
| pAG416GAL-Hsp104<sup>E190R:E191R:E192R:R419E</sup> | This paper | N/A |
| pAG416GAL-Hsp104<sup>R194E</sup> | Gates et al.<sup>59</sup> | N/A |
| pAG416GAL-Hsp104<sup>E412K</sup> | Gates et al.<sup>59</sup> | N/A |
| pAG416GAL-Hsp104<sup>R194E:E412K</sup> | This paper | N/A |
| pAG416GAL-Hsp104<sup>R353E</sup> | Gates et al.<sup>59</sup> | N/A |
| pAG416GAL-Hsp104<sup>E427K</sup> | Gates et al.<sup>59</sup> | N/A |
| pAG416GAL-Hsp104<sup>R353E:E427K</sup> | This paper | N/A |
| pAG416GAL-Hsp104<sup>R366E</sup> | Gates et al.<sup>59</sup> | N/A |
| pAG416GAL-Hsp104<sup>D434K</sup> | Gates et al.<sup>59</sup> | N/A |
| pAG416GAL-Hsp104<sup>R366E:D434K</sup> | This paper | N/A |
| pAG416GAL-Hsp104<sup>D231R</sup> | This paper | N/A |
| pAG416GAL-Hsp104<sup>D232V/R</sup> | This paper | N/A |
| pAG416GAL-Hsp104<sup>D233R</sup> | This paper | N/A |
| pAG416GAL-Hsp104<sup>K358D</sup> | Mack et al.<sup>30</sup> | N/A |
| pAG416GAL-Hsp104<sup>E360R</sup> | Tariq et al.<sup>63</sup> | N/A |
| pAG416GAL-Hsp104<sup>I361K</sup> | This paper | N/A |
| pAG416GAL-Hsp104<sup>D408K/V</sup> | This paper | N/A |
| pAG416GAL-Hsp104<sup>K410E/L/V</sup> | This paper | N/A |
| pAG416GAL-Hsp104<sup>D492R</sup> | This paper | N/A |
| pAG416GAL-Hsp104<sup>R495D/E/N/M</sup> | Tariq et al.<sup>63</sup> | N/A |
| pAG416GAL-Hsp104<sup>R496D/V</sup> | This paper | N/A |
| pAG416GAL-Hsp104<sup>D484K</sup> | Mack et al.<sup>30</sup> | N/A |
| pAG416GAL-Hsp104<sup>K480E</sup> | Tariq et al.<sup>63</sup> | N/A |
| pAG416GAL-Hsp104<sup>K480C/L</sup> | This paper | N/A |
| pAG416GAL-Hsp104<sup>K481C/E/L</sup> | This paper | N/A |
| pAG416GAL-Hsp104<sup>Y466C/K</sup> | This paper | N/A |
| pAG416GAL-Hsp104<sup>E521R/V</sup> | This paper | N/A |
| pAG416GAL-Hsp104<sup>M536K</sup> | This paper | N/A |
| pAG416GAL-Hsp104<sup>I537A/R</sup> | This paper | N/A |
| pAG416GAL-Hsp104<sup>H473F</sup> | This paper | N/A |
| pAG416GAL-Hsp104<sup>A531V</sup> | This paper | N/A |
| pAG416GAL-Hsp104<sup>R407E</sup> | This paper | N/A |
| pAG416GAL-Hsp104<sup>K470D/V</sup> | This paper | N/A |
| pAG416GAL-Hsp104<sup>K470Q</sup> | Jackrel et al.<sup>58</sup> | N/A |
| pAG416GAL-Hsp104<sup>D408K/V</sup> | This paper | N/A |
| pAG416GAL-Hsp104<sup>L414K</sup> | This paper | N/A |
| pAG416GAL-Hsp104<sup>N539L/E/D/G/K</sup> | Jackrel et al.<sup>29</sup> | N/A |
| pAG416GAL-Hsp104<sup>P411H</sup> | This paper | N/A |
| pAG416GAL-Hsp104<sup>E412K</sup> | This paper | N/A |
| pAG416GAL-Hsp104<sup>R419E</sup> | Gates et al.<sup>59</sup> | N/A |
| pNOTAG-Hsp104 | Jackrel et al.<sup>29</sup> | N/A |
| pNOTAG-Hsp104<sup>A503S</sup> | Jackrel et al.<sup>29</sup> | N/A |
| pNOTAG-Hsp104<sup>R194E</sup> | Gates et al.<sup>59</sup> | N/A |
| pNOTAG-Hsp104<sup>E412K</sup> | Gates et al.<sup>59</sup> | N/A |
| pNOTAG-Hsp104<sup>R194E:E412K</sup> | This paper | N/A |
| pNOTAG-Hsp104<sup>R353E</sup> | Gates et al.<sup>59</sup> | N/A |
| pNOTAG-Hsp104<sup>E427K</sup> | Gates et al.<sup>59</sup> | N/A |
| pNOTAG-Hsp104<sup>R353E:E427K/R</sup> | This paper | N/A |
| pNOTAG-Hsp104<sup>R366E</sup> | Gates et al.<sup>59</sup> | N/A |
| pNOTAG-Hsp104<sup>D434K</sup> | Gates et al.<sup>59</sup> | N/A |
| pNOTAG-Hsp104<sup>R366E:D434K/R</sup> | This paper | N/A |
| pNOTAG-Hsp104<sup>E360R</sup> | Tariq et al.<sup>63</sup> | N/A |
| pNOTAG-Hsp104<sup>S535E</sup> | Tariq et al.<sup>64</sup> | N/A |
| pRS313HSE-ccdB | Gates et al.<sup>59</sup> | N/A |
| pRS313HSE--Hsp104<sup>WT</sup> | Gates et al.<sup>59</sup> | N/A |
| pRS313HSE--Hsp104<sup>E190K</sup> | This paper | N/A |
| pRS313HSE--Hsp104<sup>R419E</sup> | This paper | N/A |
| pRS313HSE -Hsp104<sup>R194E:E412K/R</sup> | This paper | N/A |
| pRS313HSE -Hsp104<sup>R353E:E427K/R</sup> | This paper | N/A |
| pRS313HSE -Hsp104<sup>R366E:D434K/R</sup> | This paper | N/A |
| pTrc99A-GroEL<sub>trap</sub> | Jackrel et al.<sup>29</sup> | N/A |
| pBAD-RepA<sub>1-15</sub>-GFP | Lopez et al.<sup>79</sup> | N/A |
| pE-SUMO-Hsp72 | This study | N/A |
| pHis-TEV | Cupo and Shorter<sup>83</sup> | N/A |
| pEGFP-FUS<sup>WT</sup> | Fare et al.<sup>80</sup> | N/A |
| mCherry2-N1 | N/A | AddgeneCat#54563 |
| PMV-Hsp104-mCherry | This study | N/A |
| PMV- Hsp104<sup>A503S</sup>-mCherry | This study | N/A |
| PMV- Hsp104<sup>E191R:R419E</sup>-mCherry | This study | N/A |
| PMV- Hsp104<sup>E360R</sup>-mCherry | This study | N/A |
| PMV- Hsp104<sup>K481E</sup>-mCherry | This study | N/A |
| PMV- Hsp104<sup>E535R</sup>-mCherry | This study | N/A |
| **Software and algorithms** |  | N/A |
| Prism 9 | GraphPad |  |
| ImageJ | Rueden et al.<sup>85</sup> | N/A |
| PyMOL 4.6.0 | Schrodinger, LLC | N/A |
| Discovery Studio Visualizer | Dassault<br>Systemes Biovia<br>Corp | N/A |

## RESOURCE AVAILABILITY

### Lead contact

Further information and requests for resources and reagents should be directed to and will be fulfilled by the lead contact, James Shorter.

### Materials availability

Plasmids newly generated in this study will be made readily available to the scientific community. We will honor requests in a timely fashion. Material transfers will be made with no more restrictive terms than in the Simple Letter Agreement or the Uniform Biological Materials Transfer Agreement and without reach through requirements.

### Data and code availability

Any additional information required to reanalyze the data reported in this paper is available from the lead contact upon request.

## EXPERIMENTAL MODEL AND SUBJECT DETAILS

### Yeast strains

Yeast strains used were wild-type W303a (*MATa, can1-100, his3-11, 15, leu2-3, 112, trp1-1, ura3-1, ade2-1*) or the isogenic strain W303aΔ*hsp104*.^29^ The yeast strains W303aΔ*hsp104*- pAG303GAL-α-syn-YFP-pAG304GAL-α-syn-YFP, W303aΔ*hsp104*-pAG303GAL-FUS, and W303aΔ*hsp104*-pAG303GAL-TDP-43, have been described previously.^29,36,38^ Yeast were grown in rich medium (YPD) or in synthetic media without amino acids used for selection. 2% sugar (dextrose, raffinose, or galactose) was added to synthetic media.

### HeLa cell maintenance

Once thawed, HeLa cells were maintained in Dulbecco’s modified Eagle’s medium (DMEM) containing high glucose, supplied by Gibco. This medium was enriched with 10% fetal bovine serum (FBS) from HyClone and 1% penicillin-streptomycin solution from Gibco. The cells were incubated in a humidified incubator at 37°C at 5% (v/v) CO2. Cells that pass passage number 20 were discarded.

## METHOD DETAILS

### Site-directed mutagenesis

Mutations were introduced into Hsp104 through QuikChange site-directed mutagenesis (Agilent) and confirmed by DNA sequencing.

### Protein purification

#### Hsp104

Hsp104 proteins were purified as previously described with the following modifications.^29^ Eluate from Affi-Gel Blue Gel was equilibrated to a low-salt buffer Q (∼100mM NaCl, 20mM Tris-HCl pH 8.0, 5mM MgCl_2_, 0.5mM EDTA and 10% glycerol) and purified via ResourceQ anion exchange chromatography. Buffer Q (20mM TRIS-HCl pH 8.0, 50mM NaCl, 5mM MgCl_2_, 0.5mM EDTA, and 10% glycerol) was used as running buffer, and the protein was eluted with a linear gradient of buffer Q+ (20mM Tris-HCl pH 8.0, 1M NaCl, 5mM MgCl_2_, 0.5mM EDTA, and 10% glycerol). The eluted protein was buffer-exchanged into high-salt storage buffer (40mM HEPES- KOH pH 7.4, 500mM KCl, 20mM MgCl_2_) plus 50% glycerol and 1mM DTT and snap-frozen.

#### GroEL_trap_

pTrc99A-GroEL_trap_ was transformed into DH5α competent *E. coli* cells (Thermo Fisher). Cells were grown in 2xYT medium with appropriate antibiotics at 37°C with shaking until OD_600_ reached ∼0.4-0.6. Protein overexpression was induced with 1mM IPTG, and cells were grown at 37°C until OD_600_∼2.0. Cells were harvested by spinning (4,658g, 4°C, 15min) and pellet was resuspended in 50mM sodium phosphate buffer and centrifuged (4,658g, 4°C, 15min). The pellet fraction was resuspended in low-salt buffer (50mM Tris-HCl pH 7.5, 1mM EDTA, 1mM DTT, 50mM NaCl) and 10mg lysozyme per g cell pellet. Sample was stirred gently for 5min, lysed through sonication, and centrifuged (30,996g, 4°C, 30min). Clarified lysate was loaded onto HiTrap Q HP column (GE Healthcare) and eluted through salt gradient using low-salt buffer (as described above) and high-salt buffer (50mM Tris-HCl pH 7.5, 1mM EDTA, 1mM DTT, 500mM NaCl).^86^ Collected fractions were exchanged into the following TKME-100 buffer: 20mM Tris-HCl pH 7.5, 100mM KCl, 10mM MgCl_2_, 0.1mM EDTA, 5mM DTT, 10% glycerol, and 0.005% Triton X-100, and snap-frozen.

#### RepA_1-15_-GFP

pBAD-RepA_1-15_-GFP was transformed into BL21 (DE3)-RIL cells. Cells were inoculated in 2xYT medium with appropriate antibiotics at 37°C with shaking until OD_600_ reached ∼0.6-0.8. Protein overexpression was induced with 1mM IPTG, and cells were grown at 30°C for 4h. Cells were harvested by spinning (4,658g, 4°C, 25min) and pellet was resuspended in 40mM HEPES-KOH pH 7.4 plus 2mM 2-Mercaptoethanol (BME) and EDTA-free protease inhibitors. Cells were lysed using a sonicator and centrifuged (30,996g, 4°C, 20min). The resulting pellet was washed twice with HM buffer (40mM HEPES-KOH pH 7.4, 20mM MgCl_2_) plus 2mM BME. After each wash, cells were centrifuged (30,996g, 4°C, 20min). The pellet fraction was then resuspended in buffer containing 8M urea, 40mM Tris-HCl pH 6.8, 500mM NaCl, 10% glycerol (v/v) and agitated slowly overnight at 25°C. The solubilized pellet was then centrifuged (30,996g, 25°C, 20min) and the supernatant was collected. The supernatant fraction was incubated with Ni-NTA beads (HisPur^TM^ Ni-NTA Resin, Thermo Scientific) pre-equilibrated in buffer containing 8M urea, 40mM Tris pH 6.8, 500mM NaCl, 10% glycerol (v/v) for 2h on a spinning wheel at 25°C at the lowest speed. The Ni-NTA beads were then washed 5 times with buffer containing 8M urea, 40mM Tris-HCl pH 6.8, 500mM NaCl, 20mM imidazole, 10% glycerol (v/v) and then washed 5 times with buffer containing 8M urea, 40mM Tris-HCl pH 6.8, 500mM NaCl, 40mM imidazole, 10% glycerol (v/v). The Ni-NTA beads were then eluted with buffer containing 8M urea, 40mM Tris-HCl pH 6.8, 500mM NaCl, 500mM imidazole, 10% glycerol (v/v). The eluate was dialyzed overnight into buffer containing 40mM HEPES-KOH pH 7.4, 20mM imidazole, 150mM KCl, 2mM BME, 10% glycerol (v/v) at 4°C and re-loaded onto a Ni-NTA column (HisTrap^TM^ HP, GE Healthcare). An imidazole gradient was applied (from 20mM to 500mM) over 20CV in buffer containing 40mM HEPES-KOH pH 7.4, 20mM imidazole, 150mM KCl, 2mM BME, 10% glycerol (v/v). The purity of eluted fractions was assessed using SDS-PAGE. Collected fractions were buffer-exchanged into HKM-150 buffer (40mM HEPES-KOH pH 7.4, 150mM KCl, 20mM MgCl_2_) plus 2 mM BME and 10% glycerol (v/v) and snap-frozen.

#### Hsp70 and Hsp40

Hsc70 and DnaJA1 were from Enzo Life Sciences. Ssa1, Ydj1 and Sis1 were purified as described.^84^

#### ATPase assay

0.25µM (monomeric) Hsp104 was incubated with ATP (1mM) for 5min at 25°C in luciferase- refolding buffer (LRB150: 25mM HEPES-KOH pH 7.4, 150mM KAOc, 10mM MgAOc, 2 mM 2- Mercaptoethanol). The final reaction buffer contained < 0.3 % of HKM-500 buffer (stock of Hsp104 is >100 µM). ATPase activity was evaluated by the release of inorganic phosphate, which was measured using a malachite green phosphate detection kit (Innova Biosciences). Background hydrolysis at time zero was subtracted.

#### Luciferase disaggregation and reactivation assay

Aggregated luciferase (100nM, monomer concentration) was incubated with Hsp104 or Hsp104 variants (1µM monomer), ATP (5mM), and an ATP regeneration system (10mM creatine phosphate, 0.25µM creatine kinase) in the presence or absence of additional chaperones Hsp70 (Hsc70 or Ssa1, various concentrations as indicated in the figure) and Hsp40 (Ydj1, DnaJA1, or Sis1), for 90min at 25°C in LRB. The final reaction buffer contained less than 1% of HKM-500 buffer. After 90min, luciferase activity was measured with a luciferase assay reagent (Promega). Recovered luminescence was measured using a Tecan Infinite M1000 or Spark plate reader.

#### Luciferase refolding assay

Native luciferase (10μM) in 6M urea was incubated on ice for 5 min. The sample was then diluted to a final luciferase concentration of 1, 2, 10 or 20 nM in to LRB150 with an ATP regeneration system (10mM creatine phosphate, 0.25 µM creatine kinase), at the indicated Hsp40 (Ydj1 or Sis1) concentration. The sample was then incubated for 90min at 25°C. After 90min, luciferase activity was measured with a luciferase assay reagent (Promega). Recovered luminescence was measured using a Tecan Infinite M1000 or Spark plate reader.

To check luciferase spontaneous refolding, native luciferase (10μM) in 6M urea was incubated on ice for 5 min. The sample was then diluted to a final luciferase concentration of 1, 2, 10, or 20 nM into LRB150 with ARS. The activity of luciferase was measured at time of dilution (0 min) and after incubated for 90 min at 25°C as described above.

#### Yeast plasmids

Hsp104 variants were under control of a galactose-inducible promoter on pAG416GAL plasmids. In thermotolerance assays, Hsp104 expression was induced by 30min incubation in 37°C through a heat inducible HSE promoter on pRS313HSE plasmids.

#### Thermotolerance assay

Hsp104 variants under the HSE promoter were transformed into W303aΔ*hsp104* yeast. Yeast cultures were grown to saturation overnight at 30°C in glucose dropout media. Cultures were normalized to OD_600_ = 0.3 and grown in glucose dropout media at 30°C for at least 4h, after which the equivalent of 6 ml culture with an OD_600_ = 0.6 was grown at 37°C for 30 min (if assessing Hsp104 expression, samples would be harvested at this stage for western blot as described above). Cultures were then heat-shocked at 50°C in 1.5ml Eppendorf tubes in an Eppendorf Thermomixer for 30min and incubated on ice for 2min. Cultures were diluted appropriately, plated on glucose dropout media, and incubated at 30°C. After 2-3 days, colonies were counted using an aCOLyte colony counter and software (Synbiosis). Spotting result presented in Figure 3B was quantified using ImageJ as described in the quantification and statistical analysis section.

#### Yeast transformation and spotting assays

Plasmids containing Hsp104 variants were transformed into yeast using a standard lithium acetate and polyethylene glycol procedure.^87^ For spotting assays, yeast cultures were grown to saturation overnight at 30°C in dropout media containing raffinose. Raffinose cultures were then normalized to an OD_600_=2. Five-fold serial dilution was performed on sterile 96-well plates and spotted onto glucose and galactose plates using a 96-bolt replicator tool. Plates were grown at 30°C for 3 days and imaged at both day 2 and day 3.

#### Western blotting

For yeast Western blotting, Hsp104 variants transformed into appropriate yeast strains were grown to saturation overnight at 30°C in dropout media containing raffinose. Cultures were normalized to OD_600_ = 0.3 and grown in galactose dropout media at 30°C to induce Hsp104 and disease substrate expression (TDP-43 and FUS cultures induced for 5h). Galactose cultures were then normalized according to OD_600_ and the equivalent of 6ml culture with an OD_600_ = 0.6 were harvested by centrifugation. Media was aspirated, and the cell pellets were resuspended in 0.1M NaOH and incubated at room temperature for 5min. Cells were pelleted again by centrifugation, supernatant removed, and pellet was resuspended in 100µL 1X SDS sample buffer and boiled for 4-5min. Samples were separated via SDS-PAGE (4-20% gradient, Bio-Rad) and transferred to a PVDF membrane (Millipore) using a Trans-Blot SD Semi-Dry Transfer Cell (Bio-Rad). Membranes were blocked for at least 1h at room temperature and then incubated with primary antibodies (rabbit anti-Hsp104 polyclonal (Enzo Life Sciences); rabbit anti-FUS polyclonal (Bethyl Laboratories); rabbit anti-TDP-43 polyclonal (Proteintech); rabbit anti-GFP polyclonal (Sigma-Aldrich); mouse anti-PGK1 monoclonal (Thermo Fisher) at 4°C overnight. Membranes were washed multiple times with PBS-T, incubated with secondary antibodies (goat anti-mouse and goat anti-rabbit, LI-COR) for 1h at room temperature, and washed again multiple times with PBS-T (final wash with PBS). Membranes were imaged using a LI-COR Odyssey FC Imaging system.

#### Toxicity spotting assay

pAG416GAL plasmids containing Hsp104 variants were transformed into W303aΔ*hsp104* yeast. Yeast cultures were grown to saturation overnight at 30°C in dropout media containing raffinose. Raffinose cultures were then normalized according to OD_600_ and five-fold serial diluted. The cultures were spotted onto two sets of glucose and galactose plates using a 96-bolt replicator tool. One set of plates was grown at 30°C, and the other at 37°C, for three days and imaged subsequently at day 2 and day 3.

#### RepA_1-15_-GFP unfoldase assay

RepA_1-15_-GFP (0.7µM) was incubated with Hsp104 or Hsp104 variants (6µM, monomeric), ATP (4mM), ARS (20mM creatine phosphate, 0.06µg/µl creatine kinase). GroEL_trap_ (2.5µM tetradecamer) was included to prevent refolding of unfolded RepA_1-15_-GFP. Hsp104 variants were buffer-exchanged into TKME-100 buffer at 25°C. Reactions were assembled on ice in TKME-100 buffer plus 20µg/ml BSA. RepA_1-15_-GFP unfolding was measured by fluorescence (excitation 395nm, emission 510nm) using a Tecan Safire^2^, which was heated to 30°C prior to reading.

#### Codon-optimized Hsp104 plasmid for human cell expression

The codon optimized Hsp104 plasmid for human cell expression were purchased through Twist by two fragments with 20nt overhangs to insert into mCherry2-N1 plasmid using Gibson Assembly. The mCherry-N1 plasmid was linearized using AgeI restriction enzyme. For mCherry-tagged Hsp104 variants, mCherry is located at the C-terminal end of Hsp104. The following Hsp104 sequence was inserted on the N-terminal site of mCherry separated by a glycine-serine linker (Gly-Gly-Ser-Gly-Gly-Gly-Ser-Gly-Gly).

ATGAATGACCAGACGCAGTTCACGGAGCGCGCGCTCACCATACTCACACTTGCACAAAAACTTGCGTCTGATCACCAGCACCCGCAGCTCCAACCCATCCATATCTTGGCAGCGTTCATTGAGACTCCAGAAGACGGGTCAGTACCCTATCTGCAGAATCTGATAGAGAAGGGAAGGTATGATTACGATTTGTTTAAAAAGGTCGTTAATCGAAACTTGGTACGGATCCCCCAACAACAGCCAGCTCCGGCTGAGATAACTCCGAGTTATGCTCTCGGAAAGGTACTGCAGGATGCAGCTAAGATTCAGAAGCAGCAGAAAGATTCATTTATCGCCCAAGATCATATTCTCTTCGCTCTGTTCAACGACTCATCCATTCAACAGATCTTCAAGGAGGCTCAGGTGGACATAGAAGCTATCAAGCAGCAGGCCTTGGAGTTGCGCGGGAACACGAGAATTGATTCCCGCGGCGCAGATACTAATACACCTCTGGAATATCTTTCTAAATATGCAATAGATATGACGGAGCAGGCCAGACAGGGCAAATTGGATCCAGTGATAGGGCGAGAGGAGGAGATTCGCTCAACTATTCGAGTCCTTGCTCGAAGAATAAAAAGCAACCCATGTCTGATTGGTGAACCGGGAATTGGTAAGACTGCAATCATCGAAGGCGTTGCTCAGAGAATCATCGATGACGATGTGCCAACCATACTTCAGGGGGCGAAGCTGTTTAGTCTCGATCTTGCTGCCCTTACCGCTGGTGCAAAGTACAAAGGCGACTTTGAAGAGCGGTTTAAGGGTGTCCTCAAGGAAATCGAGGAATCAAAGACCCTTATCGTGCTTTTCATAGACGAGATTCATATGTTGATGGGGAATGGGAAAGATGATGCGGCTAACATACTCAAGCCTGCGCTCTCACGAGGACAGCTCAAGGTTATTGGCGCTACTACCAACAACGAGTACAGATCAATAGTTGAAAAGGACGGCGCGTTCGAACGGCGGTTTCAAAAAATAGAAGTAGCTGAGCCGAGCGTGAGACAGACTGTCGCCATATTGAGGGGTCTCCAGCCTAAGTACGAAATCCATCACGGCGTGCGGATCCTGGACTCAGCACTGGTTACAGCGGCGCAGTTGGCGAAACGGTATCTTCCCTACCGCAGGTTGCCCGACTCTGCTCTTGACTTGGTAGACATAAGTTGTGCGGGCGTGGCAGTTGCAAGAGACTCCAAACCTGAAGAATTGGACTCCAAAGAGCGACAACTCCAACTGATCCAGGTCGAGATTAAAGCGTTGGAGCGCGACGAAGACGCGGACTCTACTACTAAGGACCGGCTTAAACTTGCTCGACAGAAGGAAGCGTCCCTCCAGGAGGAACTCGAGCCTTTGAGGCAGCGATACAACGAGGAAAAACACGGACATGAGGAACTGACCCAAGCTAAGAAAAAGCTCGACGAGCTTGAGAACAAAGCCCTCGATGCGGAGAGGAGATATGATACTGCTACTGCTGCTGACCTGAGATACTTTGCTATCCCTGATATTAAGAAACAGATCGAAAAGCTGGAGGATCAGGTTGCTGAAGAAGAAAGACGAGCCGGAGCGAATTCAATGATACAGAACGTCGTTGATAGTGATACGATATCCGAAACAGCCGCGCGACTTACTGGAATACCGGTTAAAAAGCTCTCAGAGTCTGAGAATGAAAAACTCATTCACATGGAACGCGATCTCAGTTCAGAAGTTGTCGGTCAGATGGACGCCATTAAGGCAGTATCCAACGCTGTACGACTTTCCAGGTCTGGCCTTGCAAATCCGCGCCAACCTGCTAGCTTTCTTTTCCTTGGCCTGTCAGGGTCCGGAAAAACAGAACTGGCTAAGAAGGTTGCAGGGTTTCTGTTTAACGATGAAGATATGATGATTAGAGTAGACTGCTCTGAACTGTCCGAGAAATACGCCGTGAGTAAATTGCTCGGAACCACTGCCGGATATGTTGGATATGACGAAGGCGGATTCCTCACAAATCAGCTGCAGTACAAACCATACAGCGTCCTTTTGTTCGATGAAGTCGAGAAGGCTCACCCAGACGTTCTGACTGTTATGCTCCAGATGCTTGATGATGGGAGGATTACTTCTGGTCAAGGAAAGACCATCGATTGCAGCAACTGTATTGTAATCATGACCAGTAATTTGGGTGCTGAATTCATCAACAGTCAGCAGGGTTCAAAAATCCAAGAATCCACTAAAAACCTGGTTATGGGGGCAGTTCGGCAACACTTTCGCCCTGAATTTCTTAATCGAATCTCATCCATCGTGATATTCAACAAGCTCAGTCGCAAGGCAATCCATAAAATTGTGGACATAAGACTCAAAGAGATAGAAGAAAGGTTTGAACAGAACGATAAGCATTACAAGCTTAATCTGACACAGGAGGCAAAGGACTTCCTCGCGAAGTACGGGTATAGCGACGACATGGGTGCTAGACCGCTTAATCGCTTGATTCAAAATGAGATCCTCAACAAGCTGGCTCTTAGGATACTGAAAAACGAGATCAAGGACAAAGAGACTGTGAATGTAGTGTTGAAAAAGGGAAAATCCCGAGATGAAAATGTACCGGAAGAGGCCGAGGAATGCCTTGAAGTACTTCCAAACCATGAGGCAACCATCGGTGCTGATACCCTCGGTGATGATGATAACGAAGATTCAATGGAAATCGACGACGACCTCGAC

#### HeLa cell culture and transfections

HeLa cells were cultured in Dulbecco’s Modified Eagle Medium (DMEM) with high glucose, supplied by Gibco, which was enriched with 10% fetal bovine serum (FBS) from HyClone and 1% penicillin-streptomycin solution from Gibco. The cells were seeded into 6-well plates at a density of 2-2.5x10^5^ cells per plate, 24h prior to the transfection. The transfection was carried out with 1.5μg of total DNA mixed with 4.5μl of Lipofectamine 2000 reagent (Invitrogen). Four hours post-transfection, the medium was replaced with the standard growth medium to continue cell maintenance. 24h after the transfection, the cells were collected for analysis by microscopy or Western blotting.

In the microscopy experiments, the colocalization of proteins was assessed manually. At least 500 cells per experiment sample were analyzed across four separate trials. The statistical analysis was conducted using a one-way ANOVA with Dunnett’s test, with the calculations performed using GraphPad Prism Software.

#### Western blotting for HeLa cell experiment

For HeLa cells, ∼2-2.5x10^5^ cells were seeded and transfected with GFP-tagged FUS and either mCherry-tagged Hsp104 variants or an empty vector expressing mCherry. After 24h, cells were washed once with PBS, then resuspended in RIPA lysis buffer (150 mM NaCl, 1% Triton X-100, 1% sodium deoxycholate, 0.1% SDS, 25mM Tris–HCl pH 7.6) supplemented with protease inhibitors and 1 mM PMSF. Cells were then sonicated and centrifuged at 4°C for 10min at 10,000g, and the cell lysate was mixed with 1× SDS-PAGE sample buffer.

The samples were then boiled and separated by SDS-PAGE (4–20% gradient, Bio-Rad) and transferred to a PVDF membrane. The following primary antibodies were used: rabbit anti-GFP polyclonal (Sigma-Aldrich) for induced GFP-FUS expression, anti-alpha Tubulin monoclonal (Abcam: ab184970 for yeast; ab6160 for human cells), anti-mCherry polyclonal (Abcam). Three fluorescently labeled secondary antibodies were used: anti-rabbit (Li-Cor), anti-rat (Li-Cor), and anti-mouse (Li-Cor). Blots were imaged using a LI-COR Odyssey FC Imaging system.

#### Fluorescence microscopy

For HeLa cell microscopy, transfected HeLa cells were fixed with 2% formaldehyde for 30 min at room temperature, followed by treatment with Triton X-100 for 6 min to permeabilize cells. Coverslips were then assembled using VECTASHIELD Antifade Mounting Medium with DAPI (Vector Laboratories) and sealed before imaging. Images were taken at 100× magnification using the EVOS M5000 Imaging System (ThermoFisher) and processed using ImageJ. At least 100 cells were counted for each condition across four independent trials.

## QUANTIFICATION AND STATISTICAL ANALYSIS

The Absolute IC_50_ model in GraphPad was used to fit the dose-dependent luciferase reactivation isotherms as a function of Ssa1 or Sis1 concentrations.

Fifty=(Top+Baseline)/2

Y= Bottom + (Top-Bottom)/(1+((Top-Bottom)/(Fifty-Bottom)-1)*(AbsoluteIC_50_/X)^HillSlope)

The Bell-shaped dose-response model in GraphPad was used to fit the dose- dependent luciferase reactivation isotherms as a function of Ydj1 concentrations.

Span1=Plateau1-Dip

Span2=Plateau2-Dip

Section1=Span1/(1+(EC_50__1/X)^nH1)

Section2=Span2/(1+(X/EC_50__2)^nH2)

Y=Dip+Section1+Section2

Here, X is Ssa1, Sis1 or Ydj1 concentration, and Y is the level of reactivated Luciferase in an arbitrary unit.

Quantification is as described in the figure legends. Statistical analyses were performed using the GraphPad Prism (GraphPad Software, Inc.; La Jolla, CA, USA) as described in figure legends.

### Thermotolerance spotting assay quantification

The spotting images were opened in imageJ. The image type was changed to 8-bit, and applied background subtraction by choosing ‘subtract background’ under the ‘Process’ tab. The image was then converted to binary images by choosing ‘Binary’ under the ‘Process’ tab. The density of each spot was then quantified, as D_1_ for the first spot, and D_2_ for the second spot, etc. For each sample, only the first four spots (5-fold dilution serial) are included for this analysis. The dilution factor was corrected to account for sum of the density (D_Sum_) for each sample as shown below:

D_Sum_ = D_1_+5*D_2_ +25*D_3_+125*D_4_

The D_Sum_ of each Hsp104 variant was then normalized to D_Sum_ of Hsp104 for each replicate.

## Acknowledgements

We thank Zarin Tabassum, Zhuoyi Chen, and Sabrina Lin for feedback on the manuscript. This work was supported by an Alzheimer’s Association Research Fellowship, a Warren Alpert Foundation Distinguished Scholars Fellowship, and a Mildred Cohn Distinguished Postdoctoral Award (J.L.), a Blavatnik Family Foundation Fellowship (E.C.), and NIH grants: T32GM008076 (E.C.), R01GM110001 (D.R.S), and R01GM099836 (J.S.).

## Author contributions

Conceptualization, J.L., D.R.S. & J.S.; Methodology, J.L., E.C., & J.S.; Validation, J.L., P.J.C., C.W.G., & N.M.K.; Formal analysis, J.L; Investigation, J.L., P.J.C., C.W.G., & N.M.K.; Resources, J.L., P.J.C., C.W.G., N.M.K., E.C., S.N.G., A.L.Y., A.N.R., D.R.S., & J.S.; Data curation, J.L.; Writing–original draft, J.L., & J.S.; Writing–review and editing, J.L., P.J.C., C.W.G., N.M.K., E.C., S.N.G., A.L.Y., A.N.R., D.R.S., & J.S.; Visualization, J.L., D.R.S. & J.S.; Supervision, J.L., D.R.S., & J.S.; Project administration, J.L., D.R.S., & J.S.; Funding acquisition, J.L., E.C., D.R.S., & J.S.

---

**Figure S1.**
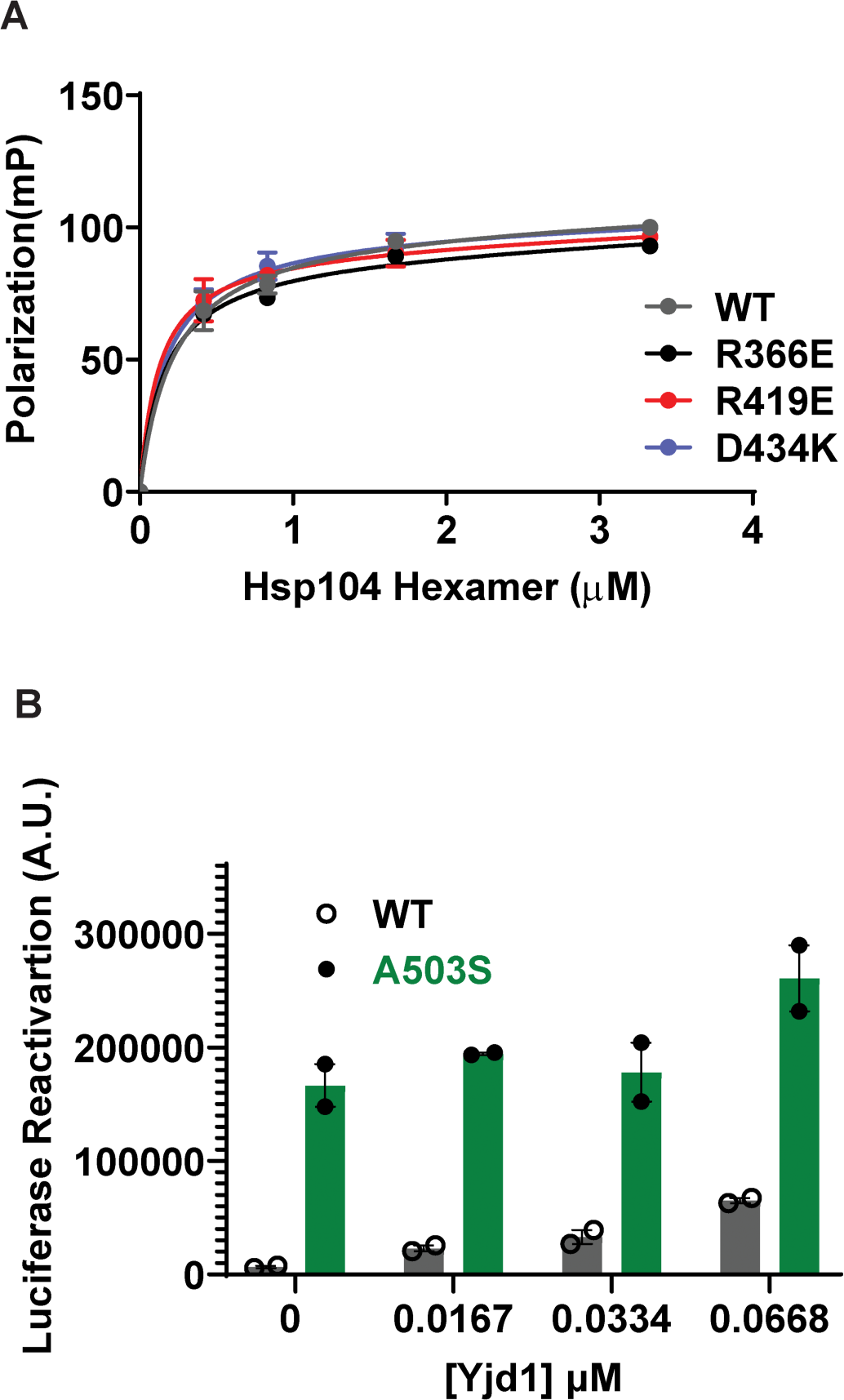
Hsp104 variants bind a model, disordered substrate, β-casein, with the same affinity as Hsp104. (**A**) FITC-casein (30nM) was incubated with the indicated concentration of Hsp104 (x-axis) in the presence of ATPγS (2mM). Binding was assessed by fluorescence polarization. Values represent means±SEM (N=2). The data were fitted using a one-site binding curve in Graphpad, and the apparent *K_D_* of the Hsp104 variants tested are similar to WT Hsp104 (0.2±0.1μM). (**B**) Bar graph of the data presented in Figure 2E for luciferase disaggregation and reactivation by Hsp104 or Hsp104^A503S^ (1µM, monomeric), plus Ssa1 (0.167µM) and the three lowest Ydj1 concentrations or in the absence of Ydj1. Bars represent means±SEM (N=2); each data point represents an independent replicate. Related to Figure 1 and 2.

**Figure S2.**
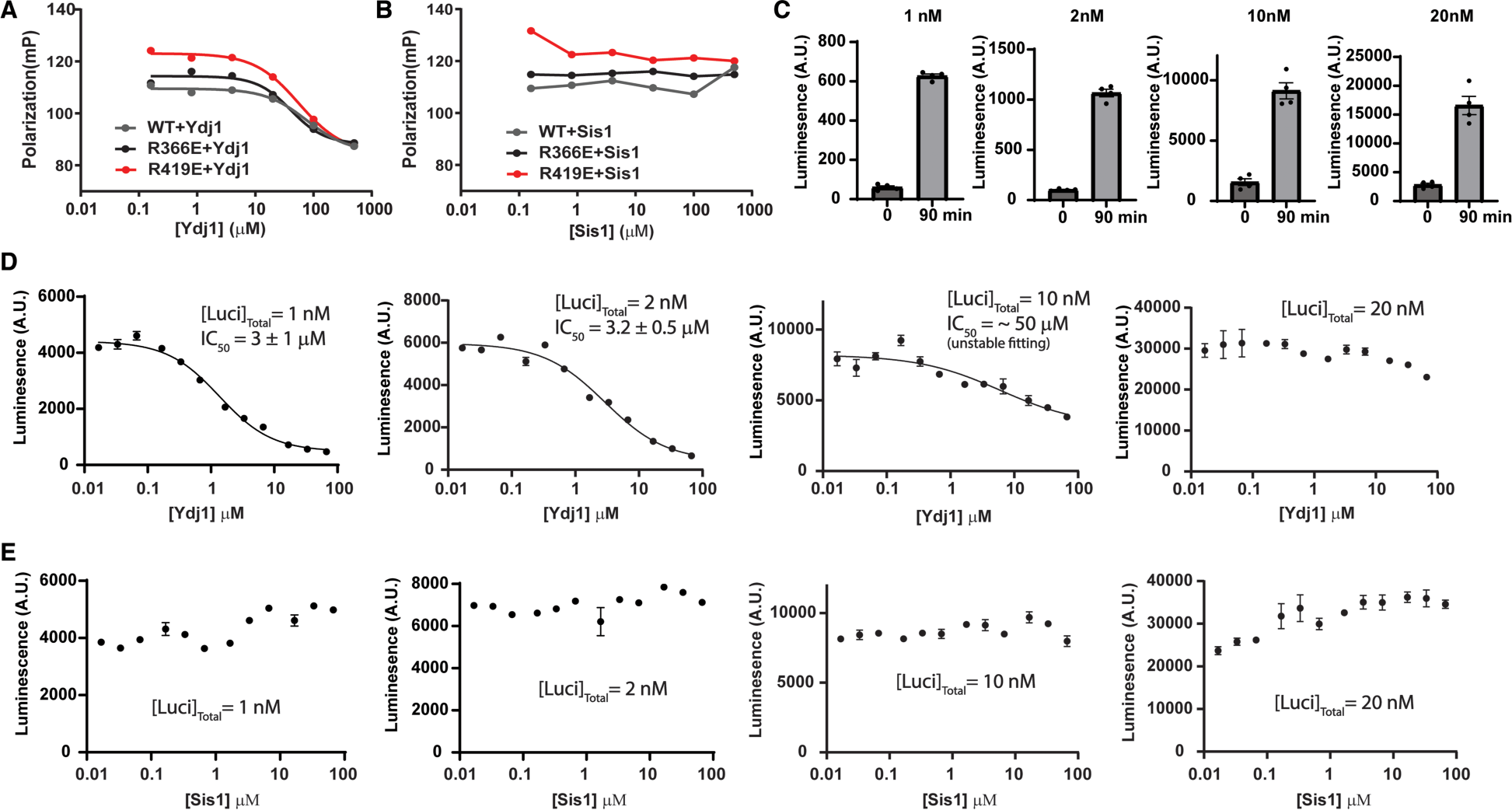
Ydj1 but not Sis1 can dissociate substrate from Hsp104 and inhibit the spontaneous refolding of unfolded luciferase. **(A, B)** Fluorescence polarization experiments measuring substrate binding competition between Hsp104 and Hsp40. Hsp104 (5μM hexameric) and the model substrate, FITC-casein (30 nM), were incubated with ATPγS (2mM) for 30min. The complex was then titrated with Ydj1 (A) or Sis1(B) at the indicated concentrations (x-axis, log scale) in the presence of ATPγS (2mM). Fluorescence polarization of FITC-casein (y-axis) was measured. Results from a representative experiment are shown. (**C**) Spontaneous refolding of soluble unfolded luciferase in buffer was measured at time of unfolding (0 min) and after 90 min. Luciferase (10µM) in 6M urea was incubated on ice for 5min and then diluted into solutions to a final concentration of 1, 2, 10 or 20nM as indicated in the figure. Luciferase activity was measured right after the unfolding reaction or after 90min in buffer. Bars represent means±SEM (N=4), each replicate is shown as a dot. (**D, E**) Luciferase (10µM) in 6M urea was incubated on ice for 5 min and then diluted into solutions containing various concentrations of Ydj1 (panel D x-axis, log scale) or Sis1 (panel E x-axis, log scale) to a final concentration of 1, 2, 10 or 20nM as indicated. Luciferase activity was measured after 90min. Values represent means±SEM (N=2). The IC_50_ of Ydj1 inhibition was determined using the dose-dependent fitting model for absolute IC_50_. Related to Figure 2 and Table S1.

**Figure S3.**
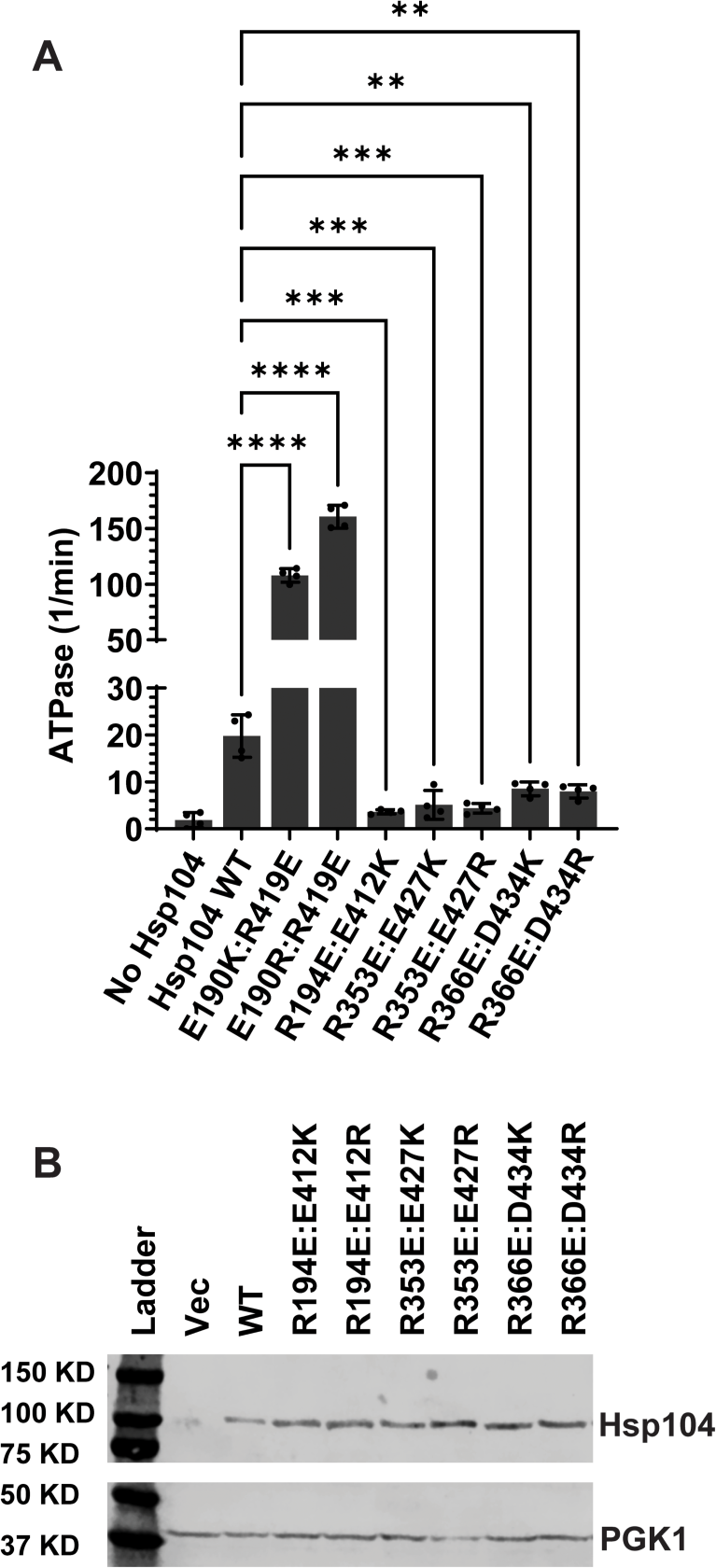
Rebuilding of the NBD1:MD salt bridges alters the ATPase activity of Hsp104. (**A**) ATPase activity of the indicated Hsp104 variants (0.25µM, monomeric) in ATP (1mM) after 5min at 25°C. Bars represent means±SEM (N=4), individual replicates are shown as dots. Dunnett’s multiple comparisons were performed to compare the ATP hydrolysis rate of NBD1- MD variants to WT. **** P≤0.0001, ***P≤0.001, **P≤0.01. (**B**) Western blots to evaluate Hsp104 expression level of yeast in the thermotolerance assay (Figure 3B). Hsp104 variants were expressed for 30min at 37°C in Δ*hsp104* yeast. Yeast were then lysed, and the lysates were processed for Western blot. PGK1 serves as a loading control. Related to Figure 3.

**Figure S4.**
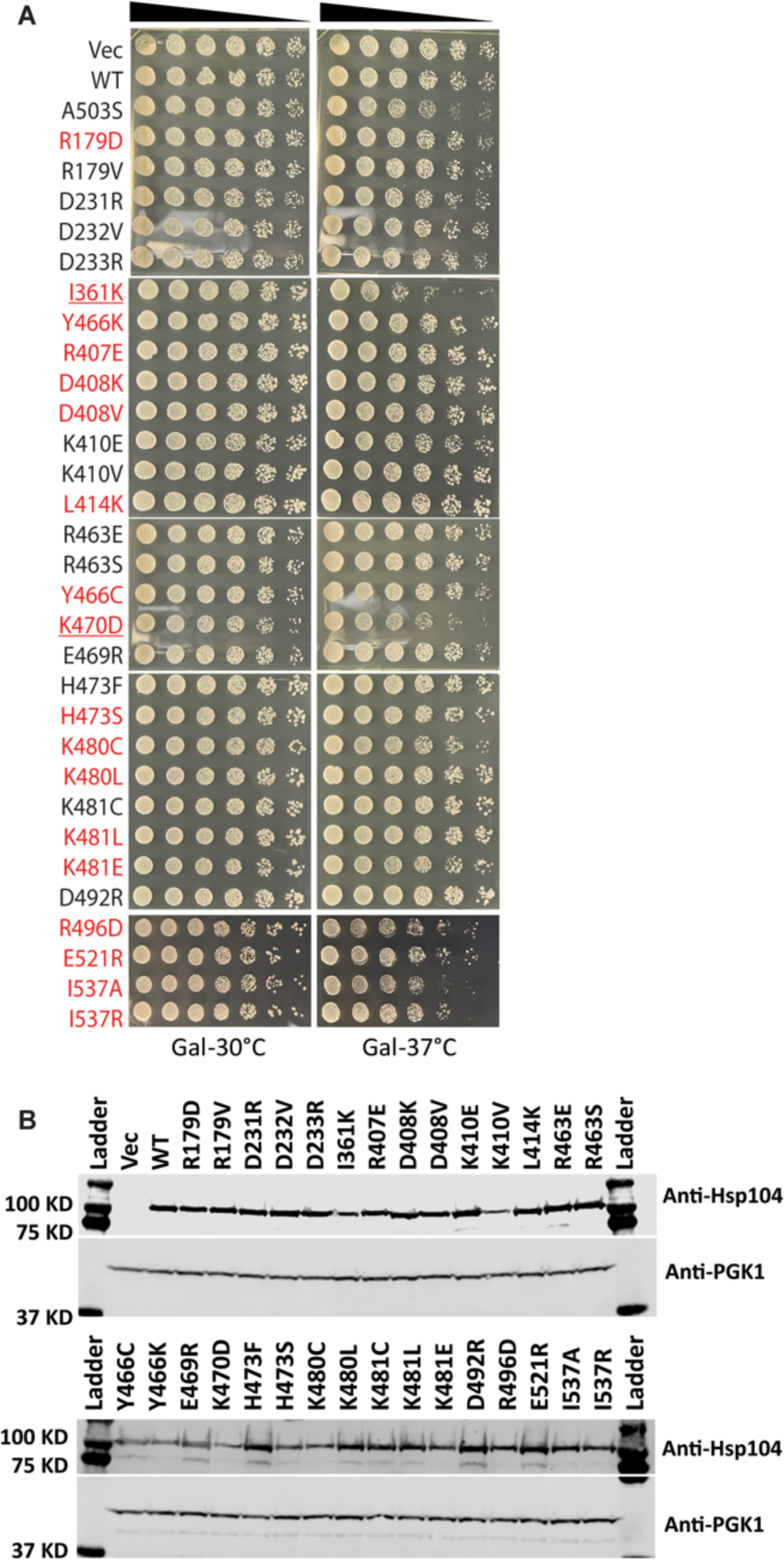
Hsp104 variant off-target toxicity and expression level in yeast. (**A**) The off-target toxicity of Hsp104 variants that perturb the intraprotomer NBD1:MD contacts of the ADP state is evaluated at 37°C using yeast spotting assay. *Δhsp104* yeast were transformed with galactose-inducible Hsp104 variants or an empty vector, WT Hs104 or Hsp104^A503S^ serve as controls. The yeast were spotted onto galactose (induction on) media in a five-fold serial dilution and incubated at 30°C (left) or 37°C (right). The potentiated variants revealed in Figure 4 are highlighted in red, and the toxic variants are underlined. (**B**) Western blots were performed to evaluate Hsp104 expression. Δ*hsp104* yeast from Figure 4 harboring the indicated Hsp104 variants or empty vector control were induced in galactose media for 5 hours at 30°C. Yeast were lysed and the lysates were visualized via Western blot. PGK1 serves as a loading control. Related to Figure 4.

**Figure S5.**
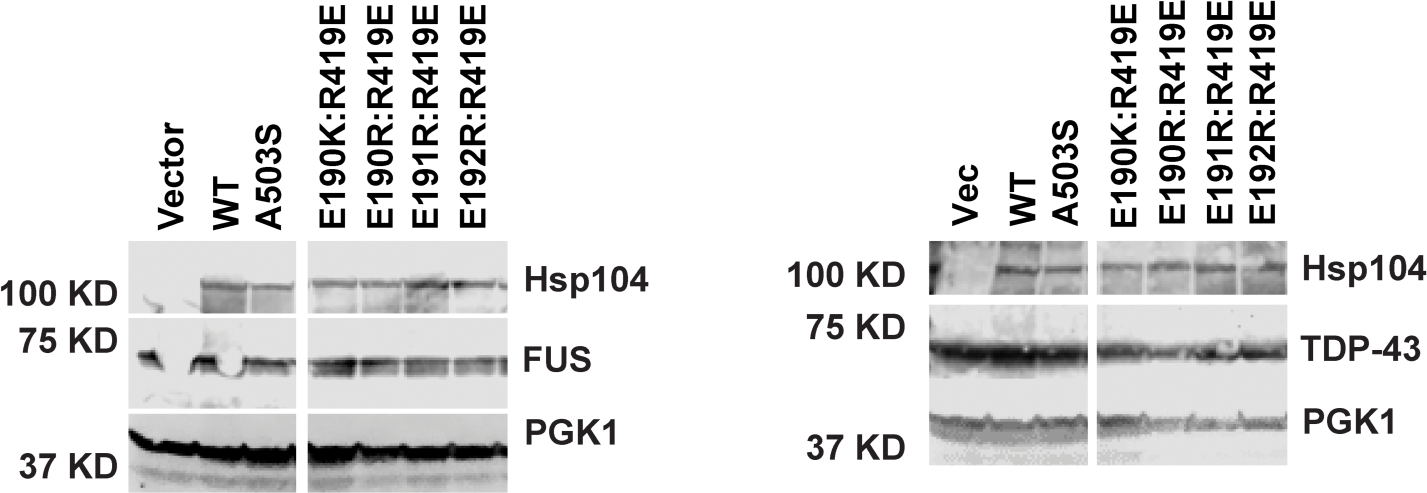
Western blots confirm Hsp104 variants and disease proteins are expressed at similar levels. Integrated Δ*hsp104* yeast strains from Figure 6C (left) and 6D (right) were induced for 5 hours in galactose media. Yeast were lysed and processed for western blot. PGK1 serves as a loading control. Related to Figure 6.

**Table S1.** Summary of EC_50_ and IC_50_ values. From left to right: 1. EC_50_ of Ssa1 for stimulation of luciferase disaggregation and reactivation by Hsp104 variants in the absence of Hsp40 (Figure 2B). 2. EC_50_ of Sis1 for stimulation of luciferase disaggregation and reactivation by Hsp104 variants in the presence of Ssa1 (Figure 2D, F). 3. EC_50_ of Ydj1 for stimulation of luciferase disaggregation and reactivation by Hsp104 variants in the presence of Ssa1 (Figure 2C, E). 4. IC_50_ of Ydj1 for stimulation of luciferase disaggregation and reactivation by Hsp104 variants in the presence of Ssa1 (Figure 2C, E). 5. IC_50_ of Ydj1 for dissociating β-casein from Hsp104 variants (Figure S2A).

|  | **1. Ssa1**<br>**EC<sub>50</sub>**<br>**(Figure 2B)** | **2. Sis1**<br>**EC<sub>50</sub> (Figure**<br>**2D, F)** | **3. Ydj1**<br>**EC<sub>50</sub>**<br>**(Figure 2C,**<br>**E)** | **4. Ydj1 IC<sub>50</sub>**<br>**(Figure 2C,**<br>**E)** | **5. Ydj1**<br>**IC<sub>50</sub>**<br>**(Figure**<br>**S2A)** |
| --- | --- | --- | --- | --- | --- |
| **Hsp104** | ~5μM | ~2μM | ~0.4μM | ~14μM | ~40μM |
| **Hsp104<sup>R366E</sup>** | ~6μM | ~1.9μM | ~0.047μM | ~0.63μM | ~20μM |
| **Hsp104<sup>R419E</sup>** | ~5μM | ~3μM | ~0.08μM | ~1.3μM | ~30μM |
| **Hsp104<sup>A503S</sup>** | ND | ~2μM | ~0.21μM | ~6μM | ND |

